# DNA double strand breaks lead to *de novo* transcription and translation of damage-induced long RNAs *in planta*

**DOI:** 10.1101/2022.05.11.491484

**Authors:** Tom Schreiber, Sunita Tripathee, Thomas Iwen, Anja Prange, Khabat Vahabi, Ramona Grützner, Claudia Horn, Sylvestre Marillonnet, Alain Tissier

## Abstract

DNA double strand breaks (DSBs) are lethal threats that need to be repaired. Although many of the proteins involved in the early steps of DSB repair have been characterized, recent reports indicate that damage induced long and small RNAs also play an important role in DSB repair. Here, using a *Nicotiana benthamiana* transgenic line originally designed as a reporter for targeted knock-ins, we show that DSBs generated by Cas9 induce the transcription of long stable RNAs (damage-induced long RNAs - dilRNAs) that are translated into proteins. Using an array of single guide RNAs we show that the initiation of transcription takes place in the vicinity of the DSB. Single strand DNA nicks are not able to induce transcription, showing that *cis* DNA damage-induced transcription is specific for DSBs. Our results support a model in which a default and early event in the processing of DSBs is transcription into RNA which, depending on the genomic and genic context, can undergo distinct fates, including translation into protein, degradation or production of small RNAs. Our results have general implications for understanding the role of transcription in the repair of DSBs and, reciprocally, reveal DSBs as yet another way to regulate gene expression.

## INTRODUCTION

DNA double strand breaks (DSBs) are severe lesions that are lethal to cells and need to be repaired correctly to prevent chromosomal rearrangements or loss of genomic information. DSBs lead to the activation of signaling cascades and repair activities called collectively the DNA-damage response (DDR), and are repaired via one of the two main repair pathways, namely non-homologous end joining (NHEJ) or homology-directed repair (HDR) (1,2). Repair via NHEJ allows only minor DNA end processing and recruits factors for direct ligation of the DSB ends leading to small insertions or deletions (indels). By contrast, HDR is accompanied by 5’-end resection leaving longer stretches of single-stranded DNA (ssDNA) 3’ overhangs that can then invade homologous sequences e.g. in sister chromatids for high-fidelity repair. Whereas NHEJ is active during the complete cell cycle, HDR is prominent in the G2/S phase, during which sister chromatids are available (1).

Another common feature of DDR is the global shut-off of canonical transcription and transcriptional arrest of genes around the DSBs, processes that are controlled via several distinct signaling pathways (3,4). In contrast to general transcriptional repression, however, transcription is also activated at the site of DNA damage, forming long divergent transcripts flanking the DSB as shown in yeast and mammalian cells (5-8). These transcripts are called damage-induced long non-coding RNAs (dilncRNAs) and there is growing evidence for their association with DSB repair in mammalian cells and yeast (5,6,9,10). It was shown that mammalian core RNA-Polymerase II (RNA-Pol II) has an intrinsic affinity for DNA ends and can use them to initiate transcription, a process that is further promoted in the presence of the Mre11, Rad50 and Nbs1 complex (MRN) (6,11). Several reports also show physical interaction between RNA Pol-II and MRN (6,11). All components of the transcription pre-initiation complex (PIC) could be detected in close proximity (i.e. within 100 bp) of the DSB whereas RNA Pol-II was located up to 2 kb downstream and upstream of the DSB (10). A current model is that RNA Pol-II is recruited to DSBs via interaction with MRN and initiates transcription from both sides of the DSB (11). According to the central role of MRN in DSB sensing and signaling for activation of repair, production of dilRNAs is likely to be an early event in DNA DSB response mammalian cells and yeast (12). Furthermore, dilncRNAs can reanneal to the DNA template strand, thereby forming DNA:RNA hybrids (short R-loops), a phenomenon that is enhanced if the DNA ends get resected from the 5’-end during HDR. In yeast and mammalian cells, DNA:RNA hybrids impact DSB repair by modulating repair factor recruitment (5,9). An equilibrium between the formation of DNA:RNA hybrids and their removal by RNase H or exosomes appears to ensure proper factor recruitment as well as formation of RPA- and Rad51-filaments at resected DNA ends needed for HDR (5,13).

Besides dilncRNAs, there is a second class of damage-induced RNAs called small damage-induced RNAs (small diRNAs in plant cells) or DNA damage response RNAs (DDRNAs in mammalian cells). These small diRNAs occur at DSBs in highly expressed repetitive loci (7,14-16). Mutational screens identified RNA-dependent RNA polymerases (RDRs), RNA processing ribonucleases (DROSHA, DICER) and diRNA-binding protein AGO2 to be involved in diRNA biogenesis or stability (7,14-17). Inhibition of RNA polymerase activity by cordycepin reduces diRNA abundance, supporting the assumption that diRNAs originate from de novo synthesized dilncRNAs (7). Although there is still a debate concerning the mechanism by which diRNAs are produced, pairing of dilncRNAs with mRNA from the same locus appears to be a prerequisite for DICER-mediated processing of the resulting double stranded RNA (6,7). However, diRNAs can also arise from exosome-mediated degradation of dilncRNAs (13). Furthermore, there is evidence that incorporation of small diRNAs primes AGO2 for sequence specific binding and the recruitment of Rad51 to the DSB site (18,19). Despite all these studies, the contribution of damage induced transcripts to DDR is not completely understood and the subject of current research efforts.

In comparison to yeast and mammalian cells, relatively less is known about these damage-induced transcripts in plants. Only small diRNAs could be identified from highly expressed transgenes (14-16). As in yeast and mammals, their role on *in planta* DSB repair pathways is still elusive. Here, we serendipitously discovered that DSBs produced by CRISPR/Cas cleavage lead to *de novo* transcription of long RNAs that can be translated if an open reading frame is in close proximity to the DSB. We could show that these long RNAs are produced from DSBs in transgenes, endogenes and also in transiently transformed T-DNAs in *Nicotiana benthamiana*. Our data uncovers a new layer of the DDR in plants, opening new perspectives for basic research but also for biotechnological applications.

## MATERIAL AND METHODS

### Bacterial and plant growth conditions

*Escherichia coli* strain DH10B [F-mcrAD(mrr-hsdRMS-mcrBC)F80dlacZDM15 DlacX74 endA1 recA1 deoR D(ara,leu)7697 araD139 GalU galK nupGrpsL l-] and *Agrobacterium tumefaciens* strain GV3101::pMP90 (Koncz and Schell, 1986) were grown in lysogeny broth/medium (LB medium [Duchefa Biochemie]: 10 g/l tryptone, 10 g/l sodium chloride, 5 g/l yeast extract) with selective antibiotics at 37°C and 28°C, respectively. *Nicotiana benthamiana* plants were grown in a phytochamber (day and night temperatures of 21°C and 18°C, respectively) with 16-h light and 50% to 60% humidity.

### Construct design and vectors

All constructs have been generated via Golden-Gate cloning (GG-cloning) with the syntax of the modular cloning system (MoClo (20,21)). Artificial PLTB-specific dTALEs were generated using the non-repetitive repeat library (22) as described in (23); RVDs: NI – Adenine, NN – Guanine, HD – Cytosine, NG - Thymine). For generation of artificial PLTB-locus specific TALEs pt57 and pt12 corresponding repeat blocks were combined with Arabidopsis Actin2 promoter (pAct2 – pAGT3459), N-terminal six His tag module (6xHis – pAGT861) and OCS terminator (tOCS – pICH41432) into T-DNA vector pICH47742 (dTALE_pt12-1_ – pAGT5954, dTALE_pt12-2_ – pAGT5955, dTALE_pt57-1_ – pAGT5956, dTALE_pt57-2_ – pAGT5957). Artificial dTALE_BETA_ was assembled similar but with the short 35S promoter (pICSL13002) instead of Actin2 promoter into T-DNA vector pICH47732 (dTALE_BETA_ – pAGT2502). All SpCas9 constructs are based on the intronized version of SpCas9 (pAGM47523; (24)). Deactivated Cas9 variant (dCas9) was generated via PCR-based mutagenesis of corresponding level -1 Modules pAGM50007 (D10A) and pAGM14696 (H841A), followed by assembly of the full length dCas9 level 0 construct (pAGM50007, pAGM50431, pAGM13474, pAGM14696, pAGM7784, pAGM50023) without the TV (6x TALE AD + VP128 – level 0 module pAGM50741; (25)) activation domain (dCas9 level 0 module without stop - pAGT6395) or with the TV activation domain (; dCas9-TV level 0 module with stop - pAGM52131). TV activation domain was synthesized by Thermo-Fisher Geneart (Regensburg). All Level 0 Cas9 modules (Cas9 (pAGM47523), dCas9 (pAGT6395) and dCas9-TV (pAGM52131)) were combined with 2xp35Sshort (pICH45089) and the omega translational enhancer (pICH41402) together with a no stop-overhang OCS terminator (tOCS - pAGT5439) for dCas9 or with the stop-overhang OCS terminator (tOCS – pICH41432) for Cas9 and dCas9-TV into T-DNA vector pICH47811 (35S:Cas9:tOCS – pAGT5997; 35S:dCas9:tOCS – pAGT7934; 35S:dCas9-TV:tOCS – pAGT5469). Intronized temperature tolerant LbCas12a (ttCas12a) Level 0 modules were generated by assembly of synthesized level -1 modules (pAGT8159, pAGT5882, pAGT5884, pAGT5886(WT) / pAGT6054(E926Q) and pAGT5888) into vector pAGM3946. The ttCas12a level 0 modules (ttCas12a – pAGT8163, deactivated dttCas12a – pAGT8173) were combined with 2xp35Sshort (pICH45089) and the omega translational enhancer (pICH41402) together with a no stop-overhang OCS terminator (tOCS - pAGT5439) for ttCas12a and dttCas12a (35S:ttCas12a:tOCS – pAGT8186; 35S:dttCas12a:tOCS – pAGT9121). Intronized version of dCas12-TV was assembled with synthesized level -1 modules (pAGT5880, pAGT5882, pAGT5884, pAGT6054 and pAGT5888) into pAGM3946 leading to a no stop level 0 dCas12a module (without the temperature tolerant mutation D156R). The dCas12a level 0 module (pAGT6056) was assembled with a short 35S promoter (pICH41388), omega translational enhancer (pAGT707), N-terminal NLS module (pAGT5892), TV-activation domain (pAGT5466) and OCS terminator (pICH41432) into T-DNA vector pICH47751 (35Ss:dCas12-TV:tOCS – pAGT5895). Cas9 single guide RNAs (sgRNAs) were designed similar as in (26), with the exception that the flip extension sgRNA scaffold together with the Arabidopsis U6-26 t67 terminator were used as template for PCR-based sgRNA amplification (27-29). Corresponding sgRNA PCR fragments were combined with the *Solanum lycopersicum* U6 promoter level 0 module (SlU6p - pAGT5824). SlU6p and AtU6-26-t67 (29) terminator were generated by PCR using pDGE412_slU6-M1E (gift from David Chiasson, LMU Munich) and Arabidopsis gDNA as template, respectively. GG-modules of Cas12a crRNAs were generated by primer extension (20 nt spacer sequence flanked by directed repeats upstream and downstream) followed by GG-cloning into pAGT6272. Corresponding crRNA modules (Cas12a) were combined with *Solanum lycopersicum* U6 promoter level 0 module (SlU6p - pAGT5824) and AtU6-26-t67 terminator level 0 module (pAGT6271) into T-DNA vector pICH47751 (crR1 – pAGT9647, crR2 – pAGT9648, crR3 – pAGT9649). Synthetic promoters for dilRNA reporter constructs (STAP, Bs4m and 35Sm) were generated as level 0 modules via PCR introducing the sgRNA B1 target site followed by GG-cloning into the corresponding vector (pAGM4023). The resulting promoter modules were combined with a GUS with introns (pICH75111) and OCS terminator (pICH4132) level 0 modules into T-DNA vector pICH47772 (B1-STAP1:GUS:tOCS – pAGT9508, B1-Bs4min:GUS:tOCS – pAGT9509, B1- 35Smin:GUS:tOCS – pAGT9507). All destination vectors allow Agrobacterium T-DNA transfer, harboring corresponding transcriptional units between a left border (LB) and right border (RB) for transfer into the plant cell.

### SpCas9 single guide RNA (sgRNA) design

PLTB-specific sgRNA (Cas9) and crRNAs (Cas12a) target sites for application in Nb were identified using the CRISPOR online tool (30).

### Agrobacterium mediated transient expression

Agrobacterium strains were grown on plates with rifampicin, gentamycin and the corresponding vector-specific antibiotic (100 µg/ml) for 1 or 2 d at 28°C. Agrobacterium strains were resuspended in Agrobacterium infiltration media (10 mM MES, 10 mM MgCl_2_, 150 mM acetosyringone) and diluted to an optical density of OD_600_ = 0.2. Agrobacterium cell suspensions were each mixed in the same proportions (v/v) and inoculated into leaves of 5 to 6 weeks old *N. benthamiana* plants with a needleless syringe. Inoculated spots were marked for harvesting.

### GUS reporter assay

At 2 or 3 days after Agrobacterium suspension inoculation, two leaf discs (diameter 0.9 mm) were harvested from three different plants (leaves) and analyzed by a fluorometric GUS assay (Kay et al., 2007). In contrast to Kay et al., 2007 the incubation time of the substrate 4-methylumbelliferone (MUG) with plant extracts was reduced to 15-30 min at 37°C. Error bars represent the standard deviation (SD) of three biological replicates from one single GUS experiment. The values displayed in GUS assays are the average of the biological replicates (each being the average of the two technical replicates). All experiments were repeated two to three times with similar results. For qualitative GUS assays, two leaf discs per inoculation spot were harvested two to three days post inoculation, incubated in X-Gluc (5-bromo-4-chloro-3-indolyl-β-D-glucuronide) staining solution for five hours at 37°C and bleached with absolute ethanol.

### Isolation of genomic DNA

Genomic DNA (gDNA) was isolated from Nb leaves using a modified CTAB method. Two to three days post Agrobacterium inoculation, two leaf discs (0.9 cm diameter) were harvested into 2 ml safe lock tubes together with 2 metal beads (2 mm diameter) and immediately transferred into liquid nitrogen. Material was disrupted using the TissueLyser II (Qiagen) for 30 sec at 30 Hz and dissolved in 300 µl CTAB buffer. After incubation for 20-40 minutes at 65°C, 200 µl chloroform was added followed by centrifugation (10 min, 18000 g). The aqueous supernatant was transferred into a new 1.5 ml tube with 200 µl of isopropanol and gently mixed by pipetting up and down several times, followed by centrifugation (10 min, 18000 g). The pellet was washed with 1 ml 70% EtOH and centrifuged again (5 min, 18000 g). The final pellet was dried for 10 minutes at 65 °C and dissolved in 200-300 µl of water (gDNA-sample).

### Identification of T-DNA insertion site flanking sequences

The T-DNA-flanking gDNA sequence for transgenic Nb pAGM26035 pt12 and pt57 PLTB-reporter lines was identified as described in (26). In brief, gDNA wasisolated using the Nucleospin Plant II kit from Macherey-Nagel. Isolated genomic DNA was G-tailed at random DNA breaks using dGTP together with terminal transferase (NEB cat M0315S) and used as template for nested PCRs using G-tail specific primer bap2pc (5’- gtccagagccgtccagcaacccccccccccccc-3’) together with T-DNA specific primer amin1 (5’- gagctcttatacagtatcctctcc-3’) (first PCR) and Bap2 (5’-gtccagagccgtccagcaac-3’) together with amin2 (5’-gaggaagggtcttgcctccgag-3’) (second PCR) for capture of RB-flanking gDNA sequences. Whole PCR products (smear) were cloned into vectors using a homology directed cloning protocol (31). Cloned inserts were sequenced with flanking vector-specific primers.

### RNA isolation and cDNA synthesis

Three days post inoculation, three leaf discs (0.9 cm diameter) from three individual plants were pooled and placed into 2 ml safe lock tubes containing 2 metal beads (2 mm diameter) followed by immediate transfer into liquid nitrogen. The material was disrupted using the TissueLyser II (Qiagen) for 30 sec at 30 Hz. RNA isolation was done using the Rneasy® Plant mini kit (Qiagen) and the RNase-free DNase set (Qiagen) according to the manufacturer’s manual. RNA was eluted in 30 µl and concentration was determined by Nanodrop (Implen, NanoPhotometer^®^ NP80). One µg of RNA together with oligo(dT) primers were used for the cDNA synthesis using the ProtoScript®II first strand cDNA synthesis kit (New England Biolabs) according to the manufacture’s protocol.

### Quantitative RT-PCR

The 5x QPCR Mix EvaGreen® (No Rox) from Bio & Sell GmbH (Germany) was used to determine the relative gene expression by real-time quantitative analysis (qRT-PCR). A master mix was created for each template cDNA and mixed with the corresponding primer pairs in the individual reactions. The amplification of the target genes was detected over 40 cycles. The data were evaluated using the CFX Maestro ™ software (BioRad). The final threshold cycle (Ct) was calculated from the average of three technical replicates. The NbUbe35 transcript was used as housekeeping gene for normalization. The relative change in gene expression was determined using the Delta-Delta-Ct method.

### Determination of transcriptional start sites by 5’ Race

1 μg of total RNA was used to synthesize cDNA with dTALE gene specific primer dTALE-GSP_R0 (cgacttgagcagcaggagatgc) using the ProtoScript®II first strand cDNA synthesis kit (New England Biolabs) according to the manufacturer’s manual. The synthesized cDNA was purified using Monarch® PCR and DNA Cleanup Kit (New England Biolabs). Poly-guanine was added at the cDNAs 3’ end using Terminal Transferase (Roche) and dGTPs. G-tailed cDNA was used as template for nested PCR using poly-G-specific primers Oligo_d(C)_Bap2_F (gtccagagccgtccagcaaccccccccccccccd) together with dTALE sequence-specific primer 5’RACE_dTALE_R1 (aaggttgtgctgctctgcgtc) (first PCR) and Anchor-specific primer Bap2-Anch-BpiI_F (ttgaagacatctcagtccagagccgtccagcaac) together with dTALE sequence-specific primer 5’RACE_dTALE_R2 (ttgaagacatctcggctggtttagctctcggtggc) (second PCR). PCR products from the nested reaction were cloned into the universal level 0 vector pAGM9121. Inserts were sequenced using vector-specific primers.

### Isolation of total RNA for RNA sequencing

Three days post inoculation, three leaf discs (0.9 diameter) were harvested from individual infiltration spots separately and placed into 2 ml safe lock tubes containing 2 metal balls (2 mm diameter) followed by immediate transfer into liquid nitrogen. The frozen tissue was disrupted using the TissueLyser II (Qiagen) for 20 sec at 30 Hz. Total RNA was isolated using the miRNeasy Micro Kit from Qiagen and the RNase-free DNase set from Qiagen according to the manufacturer’s manual. The RNA concentration was measured on a Nanodrop and integrity of the RNA was examined on a Bioanalyzer 2100 (Agilent). Five µg of RNA samples were sent to Novogene for whole transcriptome (short reads (14-42 nt) and long reads (150 nt)) sequencing.

### Processing of RNA-seq raw data

Galaxy platform (32) was used for RNAseq data handling, processing and analysis. Quality assessment of reads was carried out using FastQC v 0.11.5 (Andrews 2010, http://www.bioinformatics.babraham.ac.uk/projects/fastqc/) and MultiQC (33) before and after read cleanup. The adapter contaminations and low quality miRNA reads were removed using Cutadapt (34). Preprocessing and quantification of miRNA was carried out using MiRDeep2 (35). Small RNA reads (length of 18–28nt) were mapped to Nb genome and DSBs region using Geneious v. 7.0. Long paired-end reads (150 bp) were filtered by Cutadapt (34) and Trimmomatic v0.32 (36) to eliminate adapter contaminations and reads with low-quality. The nucleotide frequency for different position in mappings was calculated using Geneious v. 7.0 and visualized using Microsoft Excel v.13. bowtie2 (37) was used to map the trimmed long reads on the Nb genome and DSBs region and Geneious v. 7.0 was used to estimate the abundance nucleotide frequency for different position in the mappings. The raw data for RNA sequencing and sampling details are deposited at the NCBI as Bioproject SUB11197124.

## RESULTS

### A transgenic *Nicotiana benthamiana* reporter line to monitor dilRNAs

The experimental setup which led to the identification of dilRNAs was originally planned as a reporter system for homology directed repair (HDR) events *in planta*. The assay is based on a transgenic line of *Nicotiana benthamiana (Nb)* with a T-DNA designed for promoter/enhancer trap experiments (pAGM26035). The T-DNA contains a coding sequence for a designer Transcription Activator-Like Effector (dTALE_BETA_) with a 35S minimal promoter, which is not active, and three genes required for the synthesis of the red pigment betalain (Figure 1A). The genes for betalain biosynthesis (5GT; DODA1; CYP76AD1; (26)) are under the control of synthetic TALE-activated promoters (STAPs) that are specifically induced by the master regulator dTALE_BETA_ (38). Transcription of dTALE_BETA_ and subsequent betalain biosynthesis depend on the transcriptional activity coming from the genomic DNA (gDNA) flanking the T-DNA insertion site. The amplification of transcription of the betalain genes via the dTALE_BETA_ allows for the detection of low levels of transcription of *dTALE*_*BETA*_. Besides the betalain genes, a constitutively expressed kanamycin selection marker (pNOS:NptII) and a *GFP* under the control of a STAP, are also present on the T-DNA (Figure 1A). We selected seven primary transformants (pt) that did not show any betalain production (Supplemental Figure 1 and 2). First, we confirmed the presence of functional betalain biosynthesis genes by transient expression of dTALE_BETA_ in leaves of these transformants (Figure S1A and B). Four out of seven *Nb* transformants contain functional betalain biosynthesis genes and for two of them we could also identify the T-DNA flanking gDNA sequence (pt12 and pt57; Figure S1). We confirmed the functionality of the transgenic *dTALE*_*BETA*_ using transcriptional activators that bind in the T-DNA flanking gDNA upstream of *dTALE*_*BETA*_. These were either other dTALEs (pt12/pt57) or Cas9-based dCas9-TV activator (25) together with specific single guide RNAs (sgRNAs G1-G3) (Figure S2). Due to higher inducible expression levels, the progeny of the transgenic pt57 reporter line was used for further experiments and we refer to the transgene as the promoter-less TALE-betalain (PLTB)-locus.

**Figure 1.**
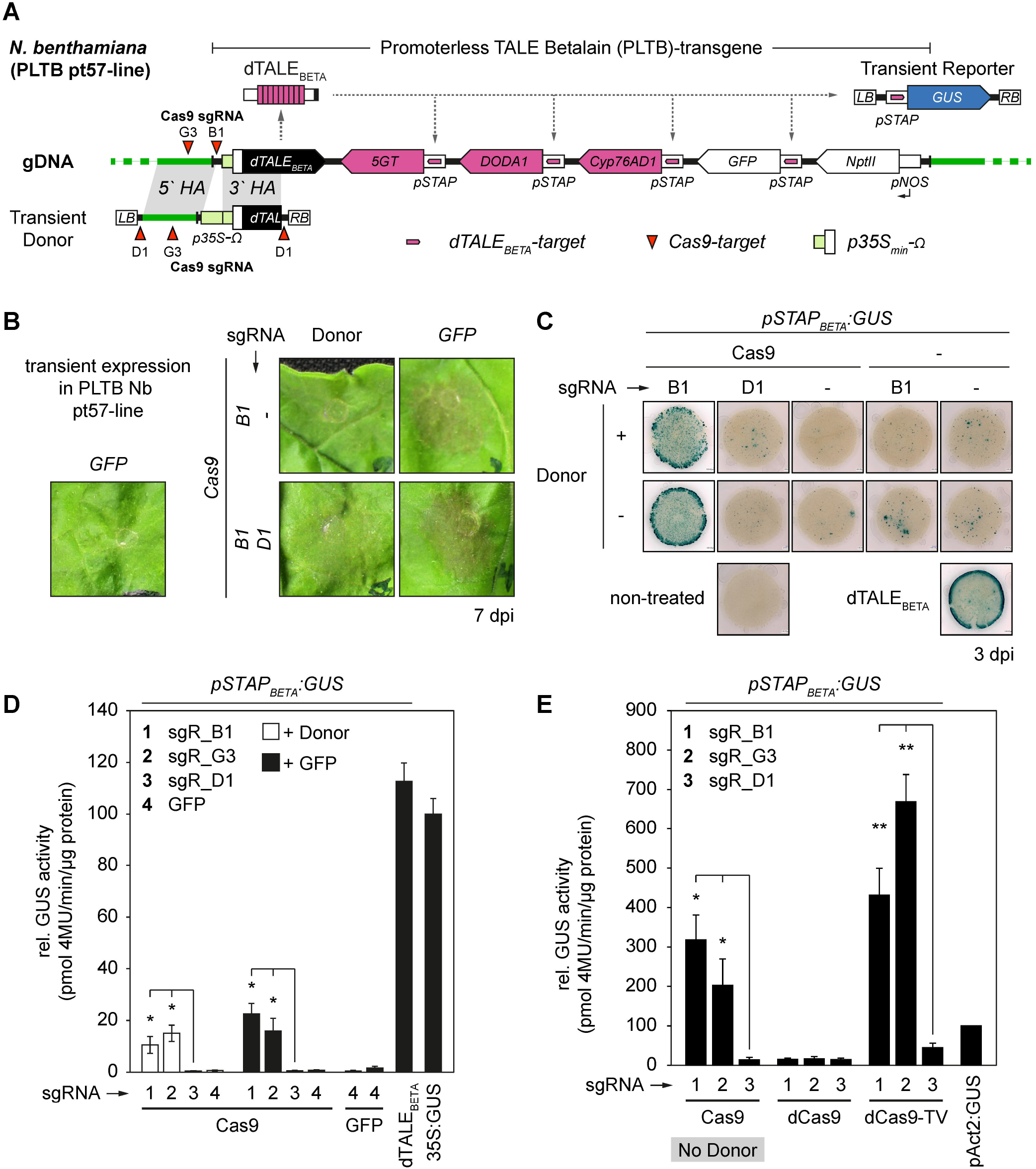
DSBs lead to de novo transcription and translation of dilRNAs at the transgenic promoter-less TALE Betalain (PLTB) locus. **(A)** Schematic overview of the PLTB transgene in *Nicotiana benthamiana* (pAGM26035 pt57). The inactive master regulator dTALE_BETA_ is supposed to be activated by recombination of the full length 35S promoter from a transiently expressed donor T-DNA using HDR of the sgRNA B1-induced lesion. Recombination of the 35S promoter upstream of dTALE_BETA_ should activate its expression and consequently activate Betalain biosynthesis (5GT, DODA1, Cyp76AD1) or GUS if the transient GUS reporter is co-expressed. **(B)** Leaf phenotypes, seven days post inoculation (7dpi) of Agrobacterium strains carrying corresponding gene targeting constructs (Cas9 and sgRNAs in the presence or absence of the 35S promoter donor). Betalain biosynthesis (red coloration) is independent of the donor and requires Cas9-mediated cleavage upstream of dTALE_BETA_ (sgRNA B1). Transient expression of GFP (35S:GFP) serves as negative control and sgRNA D1 only cleaves in the donor T-DNA. **(C)** GUS-stained leaf discs harvested 3dpi after transient expression similar to B but with co-expressed GUS reporter. Measured is the dTALE_BETA_-induced GUS activity. Untreated leaf and Transient expression of dTALE_BETA_ serves as negative and positive control, respectively. **(D and E)** Quantitative GUS measurement 3dpi after transient expression of indicated constructs. Both sgRNAs leading to cleavage upstream of dTALE_BETA_ (sgRNA B1 and G3) induce dTALE_BETA_ expression. Deactivated Cas9 (dCas9) and dCas9-TV transcriptional activator serve as controls. Transient expression of GFP and dTALE_BETA_ serve as negative and positive control, respectively. (Student’s t-test; *P-value ≤ 0.05; **P-value ≤ 0.01; ***P-value ≤ 0.001)

### DNA double strand breaks induce *de novo* transcription of messenger RNAs in planta

Initially, we used the transgenic PLTB pt57 line for a gene targeting approach, in which we introduce a DSB upstream of the master regulator *dTALE*_*BETA*_ (sgRNA B1) and aim to repair the lesion by integration of a 35S promoter via HDR (Figure 1A). Cells where a successful integration by HDR happens should produce betalain and therefore become red. We transiently expressed Cas9 (SpCas9) together with the PLTB-specific sgRNA B1 and a donor T-DNA harboring the 35S promoter (p35S) flanked by approximately 1 kb long 5’- and 3’-homology arms (HA) specific for the PLTB-locus and monitored betalain accumulation over seven days (Figure 1B). A donor-specific sgRNA D1 was also added to release the donor fragment from the T-DNA, with the aim to potentially increase HDR efficiency as shown previously (39). Expression of GFP (*35S:GFP*) alone served as negative control to confirm that the pt57 line does not produce any betalain upon Agrobacterium-mediated transient expression (Figure 1B). Surprisingly, we observed a slight red coloration (betalain accumulation) in leaf areas that express Cas9 together with sgRNA B1 but in the absence of the donor with p35S (Figure 1B). Co-expression of the donor-specific sgRNA D1 had no effect. The presence of a weak homogeneous red coloration in the infiltrated leaves in the absence of the donor DNA suggests that there is transcriptional induction of the *dTALE*_*BETA*_ located on the PLTB-transgene. In order to quantify the dTALE_BETA_ activity with increased sensitivity we repeated this experiment and co-expressed a transient GUS reporter *(pSTAP*_*BETA*_*:GUS*) which can also be induced by dTALE_BETA_ (Figure 1C to E). Here we could confirm that Cas9-mediated DNA cleavage upstream of the master regulator dTALE_BETA_ alone is sufficient to activate its expression (Figure 1C). The GUS activity induced by the dTALE_BETA_ upon Cas9 cleavage is about 20% of the GUS activity driven by the strong constitutive 35S promoter. Co-expression of Cas9 together with the control sgRNA D1, or co-expression of dCas9 together with the PLTB-specific sgRNAs B1 or G3 did not induce dTALE_BETA_ activity, demonstrating that dTALE_BETA_ expression strictly depends on cleavage by Cas9 upstream of the *dTALE*_*BETA*_ locus (Figure 1D-E). We concluded that like yeast, fungi (*Neurospora crassa*), drosophila and mammalian cells, plants also produce long DNA-damage induced transcripts (8). However, based on our observations these transcripts can be translated into proteins and we propose to call them damage-induced long RNAs (dilRNAs).

Next, we wondered whether we could also detect dilRNAs when the DSB occurs in the vicinity of an endogene. As a potential target we chose the basic helix-loop-helix transcription factor *NbUPA20* (Figure 2). *NbUPA20* is transcribed at very low levels and can be strongly induced by the natural TALE AvrBs3, which served as positive control for transcriptional induction (40). We designed three *NbUPA20*-specific sgRNAs targeting the promoter region, Figure 2A). We transiently co-expressed Cas9 or dCas9 together with corresponding *NbUPA20* sgRNAs in wild type plants and performed qRT-PCR with *NbUPA20* specific primers on cDNA generated using oligo-dT. Similar to the observations on the PLTB-locus in the PLTB pt57 line, co-expression of *NbUPA20*-specific sgRNAs together with Cas9 but not with dCas9 led to a 2.5-3.5 fold induction of *NbUPA20* transcript abundance (Figure 2B). Transient expression of dAvrBs3 led to 134 fold induction of NbUPA20 transcripts, which suggests that the abundance of dilRNAs is comparatively low. We also observed a reduced *NbUPA20* transcript abundance if dCas9 is co-expressed with sgRNA_UPA20-3 (compared to dCas9 with no sgRNA). This reduction is likely due to dCas9 interference with the transcription initiation complex that assembles close to the sgRNA_UPA20-3 binding site (CRISPR interference).

**Figure 2.**
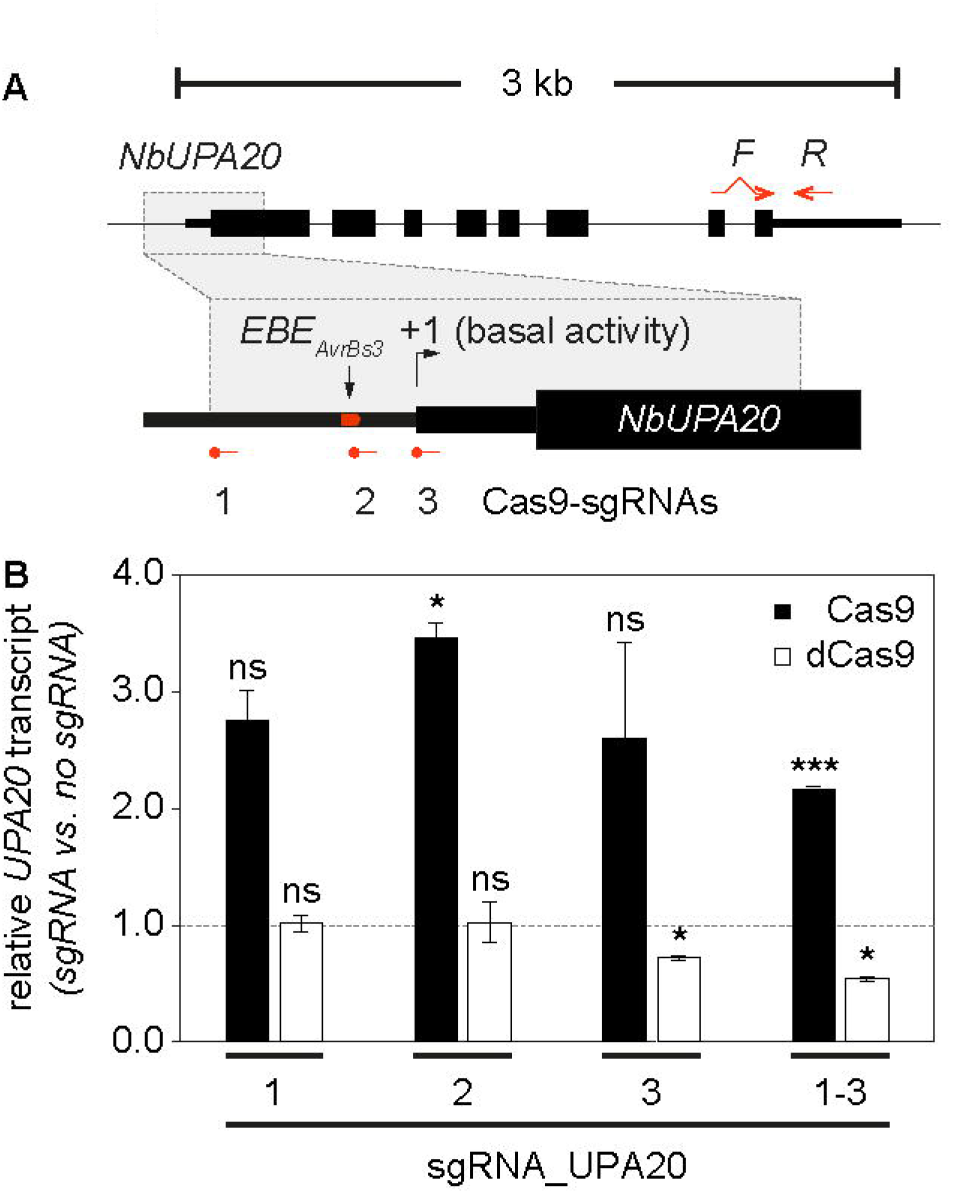
DSBs upstream of the *Nicotiana benthamiana* endogene UPA20 leads to de novo transcription of dilRNAs. **(A)** Schematic overview of the UPA20 gene organization and corresponding Cas9 (sgRNA) and dTALE AvrBs3 (effector binding element – EBE) targets. NbUPA20 is induced by AvrBs3 which serves as a positive control for transcriptional upregulation. Intron spanning primers for qRT-PCR are indicated by red arrows. **(B)** Relative NbUPA20 transcripts (sgRNA vs. no sgRNA) determined by qRT-PCR. Corresponding sgRNAs are expressed together with Cas9 (black bars) or deactivated dCas9 (white bars). DAvrBs3-mediated upregulation of NbUPA20 (relative to expressed 35S:GFP) is about 134 fold and not shown in the graph. Indicated significant changes are relative to NbUPA20 transcript without sgRNA (Student’s t-test; *P-value ≤ 0.05; **P-value ≤ 0.01; ***P-value ≤ 0.001).

In summary, we showed that DSBs lead to the activation of on target *de novo* transcription of dilRNAs in plants which can be translated into proteins.

### Blunt- and sticky-ended DSBs but not ssDNA nicks lead to translatable dilRNAs

The detection of dilRNAs in planta with our reporter system raised further questions, such as whether DNA cleavage always leads to dilRNAs and whether the site of DSB defines their transcriptional start site (TSS). To answer these questions, we first expanded the set of sgRNAs specific for the PLTB-locus by selecting positions in the genomic DNA flanking the T-DNA (G1-G3), in the 35S minimal promoter, in the omega translational enhancer (B1-B4) and in the N-terminal coding sequence of dTALE_BETA_ (T1-T6) (Figure 3A). To test a possible effect of the orientation of the sgRNA, we also designed them with the protospacer adjacent motif (PAM) in the forward or reverse configuration. In a first screen using the PLTB-specific sgRNAs, 12 out of 14 led to detectable dTALE_BETA_-induced GUS activity, which implies production of a translated dilRNA (Figure 3B). As seen before, increased GUS activity depends on Cas9-mediated DNA cleavage, because no increase could be observed if PLTB-specific sgRNAs are combined with dCas9. The dTALE_BETA_-induced GUS activities range from 50% to 300% of the pAct2-driven GUS activity and reached the highest activities if DNA cleavage occurs closely upstream of or inside the 35S minimal promoter (sgRNAs B1-B3; Figure 3B). SgRNA B4 cleaves two bases upstream of the ATG of the full length dTALE_BETA_ ORF (see annotated PLTB locus in supporting information). The lack of GUS activity in Cas9 sgRNA B4 samples suggests that two bases is too short to allow initiation of translation at this particular <u>A</u>TG_1_. The out of frame ORF (<u>A</u>TG_65_) upstream of the <u>A</u>TG_145_ (Figure 3A, black line) could prevent dTALE_BETA_ translation from <u>A</u>TG_145_. In the case of sgRNA T1, we could show that this sgRNA does not allow efficient cleavage by Cas9 (Figure S3), thus explaining the absence of dilRNAs and dTALE_BETA_ activity. N-terminal truncations in the TALE proteins lead to either no loss of transcription activation (63 amino acid deletion) or a five-fold reduction (152 amino acid deletion) (41). Therefore, a DNA cleavage downstream of the first <u>A</u>TG_1_ of dTALE_BETA_ (by sgRNAs T1-T6) could still lead to a functional TALE protein if the translation is initiated at one of the two downstream in frame ATGs at positions <u>A</u>TG_145_ (ΔN49aa) and <u>A</u>TG_361_ (ΔN121aa) respectively (Figure 3A, see indicated ORFs).

**Figure 3.**
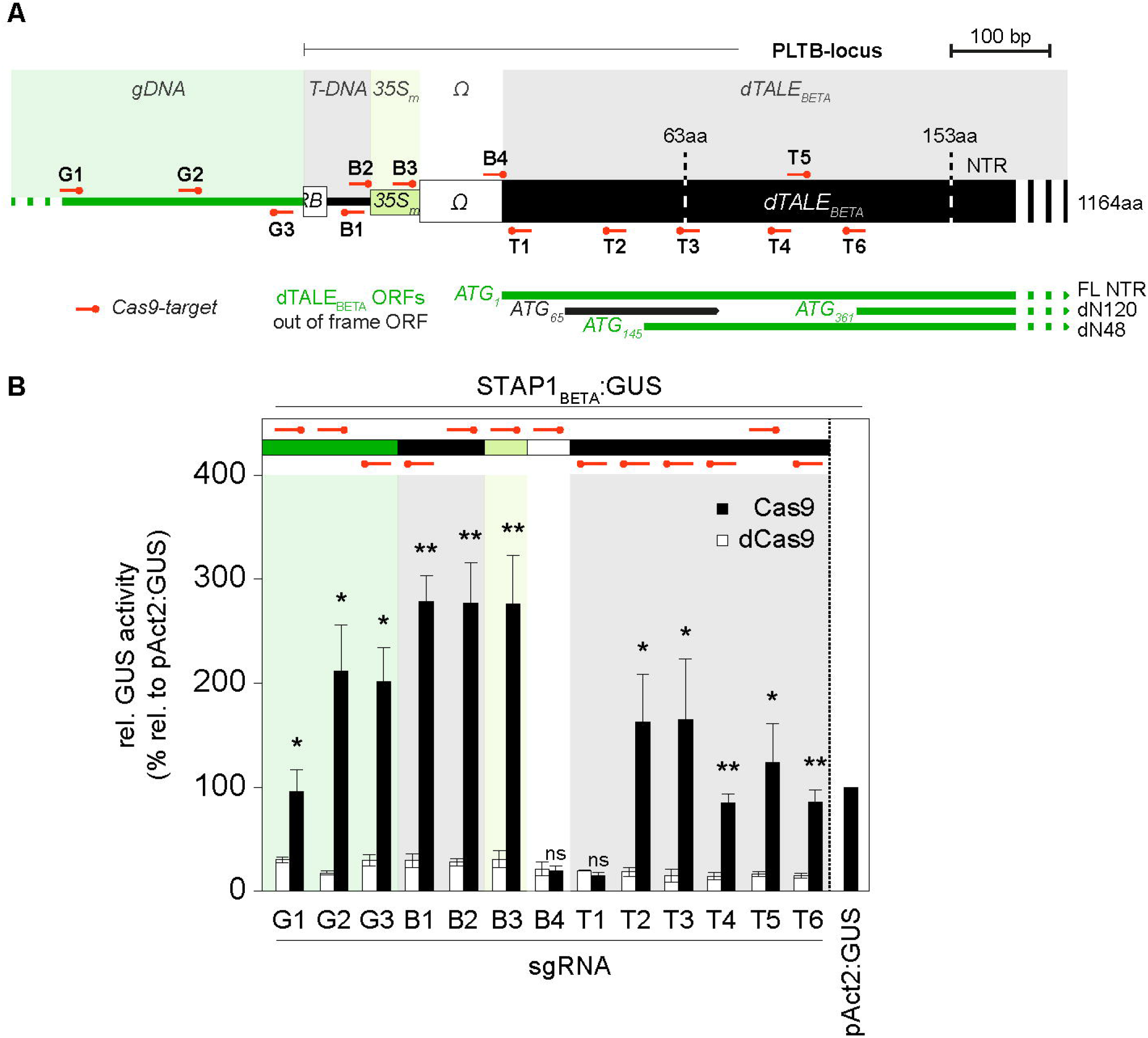
DSBs in the flanking gDNA, promoter and coding sequence of dTALE_BETA_ lead to transcription and translation of dilRNAs. **(A)** Schematic overview of tested Cas9 target sites in the PLTB-locus. Cas9 targets are localized to the T-DNA flanking gDNA (G1- G3), the transgenic promoter region (B1-B4) and the dTALE_BETA_ coding sequence (T1-T6). N-terminal deletions of dTALEs with WT-activity (ΔN63) or 5-fold reduced activity (ΔN153aa) are indicated by dashed lines. Alternative open reading frames (ORFs) within the coding sequence of dTALE_BETA_ are indicated in black (non-functional) and green (functional) with nucleotide position of the ATG and the corresponding N-terminal amino acid truncations given on the right. **(B)** Quantitative GUS measurement 3dpi after transient expression of individual sgRNAs together with Cas9 (black bars) and dCas9 (white bars). Transient expression of Actin2 promoter-driven GUS serves as positive control. (Student’s t-test; *P-value ≤ 0.05; **P-value ≤ 0.01; ***P-value ≤ 0.001)

Next, we asked whether single strand DNA nicks also induce transcription. We used nickase versions of Cas9 (D10A or H840A), which cleave only one strand, and two overlapping gRNAs separately (B1 and B2). No GUS activity could be detected, showing that nicks do not activate transcription of dilRNAs (Figure S4B). We also applied Cas9 nickases in a dual sgRNA strategy to generate DSBs with 68 nt long 5’ or 3’ overhangs. Here, no significant induction of GUS activity could be detected either (Figure S4C). We cannot exclude that nicks on opposing strands separated by 68 nucleotides might not lead to DSBs with long overhangs. To analyze whether dilRNAs are also produced from sticky ends we used Cas12a to induce DSBs with short (7nt) 5’ overhangs and could observe induction of dilRNAs to a level similar to that produced by Cas9 DNA cleavage (Figure S5). In summary, a DSB is needed for the induction of dilRNAs.

To get further insights about dilRNA-abundance we performed GUS assay and qRT-PCR for the dTALE_BETA_ in parallel (Figure 4B and Figure S6). We opted for a subset of sgRNAs (G2-G3, B1-B4, T1-T2), including the two sgRNAs which did not lead to increased GUS activity (sgRNA B4 and T1). As positive controls, we expressed in *trans* the transcriptional activators dCas9-TV with sgRNAs B1 or B2, and dTALE-pt57-2, which binds to a sequence in genomic DNA just upstream of the T-DNA (Figure 4A). As observed previously, we did not measure increased GUS activity when Cas9 is combined with sgRNAs B4 and T1 (Figure S6). We observed much stronger GUS activity induced by the activators expressed in *trans* (dCas9-TV and dTALE-pt57-2) than with DSB-induced transcripts. Since we cannot distinguish between transcripts from dTALE_BETA_ and dTALE-pt57-2, we did not include this sample in the qRT-PCR experiment (Figure 4B). Quantitative RT-PCR revealed that the abundance of dilRNAs is very low compared to transcripts induced by dCas9-TV (3% for sgRNA B1 and 5% for sgRNA B2; Figure 4B). In contrast to the GUS assay, we could detect significant amounts of dilRNAs by qRT-PCR when Cas9 is expressed with sgRNA B4, which further supports the overlapping out of frame ORF as the reason for the absence of dTALE_BETA_ translation (Figure 4B and Figure S6).

**Figure 4.**
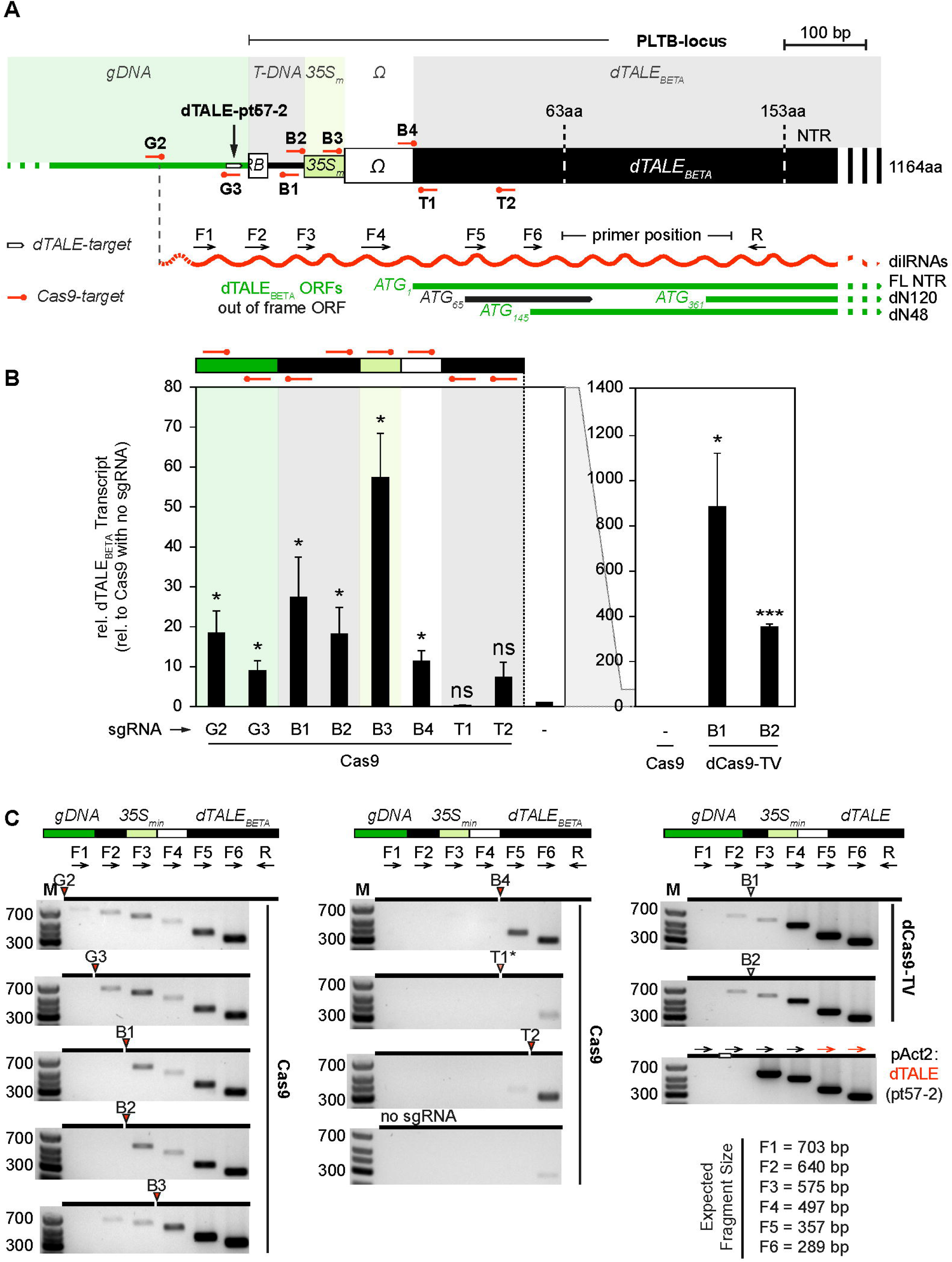
Transcriptional start sites of dilRNAs are defined by the site of DSB. **(A)** Schematic overview of tested Cas9 target sites and used primers for RT-PCR to determine the transcriptional start site of dilRNAs. Labelling is similar to Figure 3. Results from B and C came from the same samples and have been generated in parallel. **(B)** Relative dTALE_BETA_ transcripts normalized to Cas9 without sgRNA. Due to the wide range the diagram has been separated into dilRNAs and dCas9-TV induced transcripts. Transcripts induced by dTALE(pt57-2) have not been included into the graph because with the used primers we could not differentiate between dTALE_BETA_ and dTALE(pt57-2). (Student’s t-test; *P-value ≤ 0.05; **P-value ≤ 0.01; ***P-value ≤ 0.001) **(D)** Agarose Gel pictures of PCR products using indicated primers and individual cDNAs. Please note that we could not detect cleavage activity for sgRNA T1*. For cDNA from dTALE(pt57-2) induced dTALE_BETA_, primer pairs F5+R and F6+R could not distinguish between dTALE(pt57-2) and dTALE_BETA_ transcripts and are therefore indicated in red. Expected fragment sizes are given on the lower right panel.

### DSB sites define the transcription start site of dilRNAs

In order to identify the TSS of dilRNAs and activator-induced RNAs we performed 5’ RACE (Rapid Amplification of cDNA Ends) using oligo-dT or gene specific primers for the reverse transcription. We could capture PLTB-specific transcripts with the transcriptional activators (dCas9-TV and dTALE pt57-2) but not with dilRNAs most likely due to low RNA levels (Figure S7A). Interestingly, we observed that transcripts induced by dCas9-TV start at several positions and quite far downstream from the protospacer sequence, whereas in the case of dTALE pt57 most of the transcripts started within 100 bp from the binding site (Figure S7B). This information is particularly relevant for the application of dCas9-TV in engineering endeavors.

Because we could not generate sufficient product with 5’-RACE for dilRNAs we performed a RT-PCR amplification using forward primers located at different positions, thus allowing us to map the position of the TSS. We chose six primers (F1-F6) specific for the PLTB-locus that bind in between the individual Cas9 cleavage sites (Figure 4A and C; Figure S8A). With the exception of sgRNA B3 and T1 we could show that the TSS of the dilRNAs follow the site of DNA cleavage. Interestingly, combination of Cas9 with sgRNA B3 also led to transcripts that span the DSB site. Putative sgRNA B3 off-targets upstream of the on-target could not be found, indicating that this DSB-spanning dilRNAs are not induced by a DSB located upstream of the sgRNA B3 binding site. RT-PCR performed with cDNA from activator-induced transcripts revealed that in contrast to dTALEs, dCas9-TV also induces transcription upstream of its cognate binding site, although the RNAs initiated upstream of the target sequence are minor products (Figure 4C).

In summary, DSBs in plants activate the production of dilRNAs, most likely initiated directly at the DNA DSB end. Furthermore, production of dilRNAs seems to be a systematic consequence of DBSs rather than an exception, because all sgRNAs that mediated DNA-cleavage also showed formation of dilRNAs.

### dilRNAs are produced in both directions of the DSB with context dependent stability

Next, we investigated whether dilRNAs are produced in both directions as described for yeast and mammalian cells (5,6). In plants, only the production of small diRNAs upon DNA cleavage in highly expressed transgenes was identified (14,16). In addition, we wanted to know whether small diRNAs are produced besides dilRNAs after DNA cleavage. To address this, we conducted an RNA-seq experiment covering all possible RNA populations (mRNA / long non-coding RNAs (150 nt reads) and siRNAs (12-42 nt reads)). We used sgRNA B1 together with Cas9 and the transient GUS reporter to confirm the production of dilRNAs (Figure S9). Expression of the sgRNA B1 without Cas9 served as negative control. We could detect long RNA reads on both sides of the DNA after DSB, but dilRNAs downstream of the cleavage site are significantly more abundant. This likely reflects stabilization of the RNA via polyadenylation thanks to the presence of the terminator downstream of the *dTALE*_*BETA*_ (Figure 5A). The absence of a terminator and of an ORF in the sequence upstream of the cleavage site might promote dilRNA degradation by nonsense-mediated decay. By contrast, diRNAs were hardly detected in the RNA-seq of small reads (diRNAs) (Figure 5B). Furthermore, we could map several small RNA reads in the vicinity of the cleavage site. However, sequence analysis revealed that these reads derive from the constructs that are expressed transiently (pAGT5997: <u>35S:Ω</u>-Cas9; pAGT1963: STAP:<u>Ω</u>-GUS_pNOS-<u>Ω</u>-BAR-tNOS and pAGT6149: U6 sgRNA <u>B1</u>, origin underlined). The 35S promoter peak is absent in the control without Cas9. Small RNA reads derived from sgRNA B1 are also detected in the negative control but with lower abundance. The GUS reporter also contains a constitutively expressed BAR resistance gene on the same T-DNA which is likely the origin of most small reads mapped to the omega enhancer sequence (Figure S10). This explains the presence of small RNAs even in the absence of Cas9. The low number of small RNA reads mapping to the genomic sequence upstream of the cleavage site are present independently of Cas9 and probably originated from homologous sequences, as seen by mismatches in the mapped sequence. In summary, we could detect dilRNAs and also confirmed that these transcripts are terminated and polyadenylated if a terminator is present. Furthermore we could not detect significant amounts of diRNAs in response to DNA cleavage within the silent PLTB-locus.

**Figure 5.**
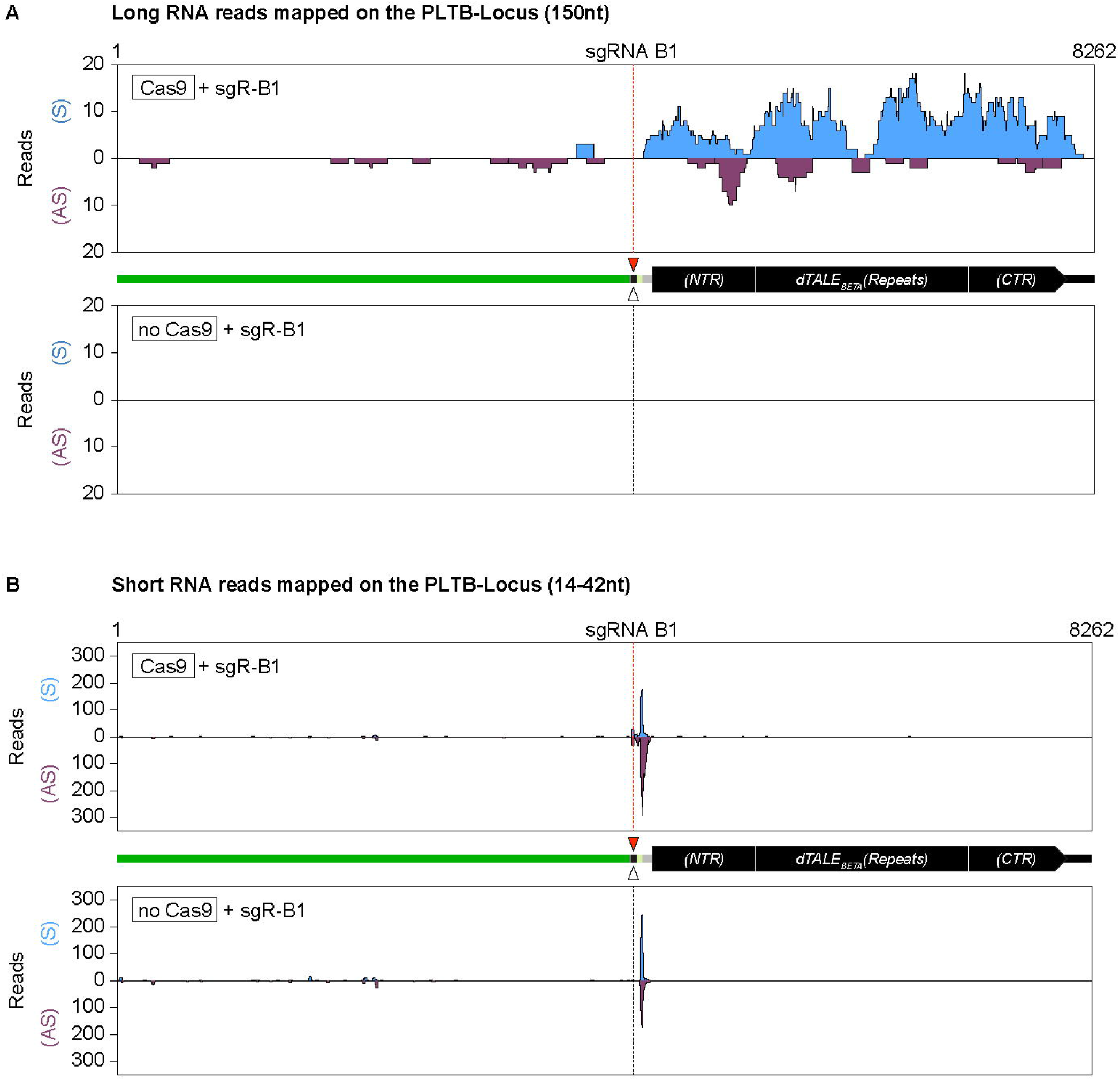
Divergent production of dilRNAs upstream and downstream of DSBs. **(A)** Mapping of long (150 nt) RNA-seq reads to the PLTB-locus. The upper and the lower panel shows long RNA reads derived from samples with and without Cas9 expression, respectively. **(B)** Mapping of short (14-42 nt) RNA-seq reads to the PLTB-locus. The upper and the lower panel shows short RNA reads derived from samples with and without Cas9 expression, respectively.

### Induction of dilRNAs also occurs on transiently expressed T-DNA

We wondered whether the dilRNAs induced by DSBs in genomic DNA could also be induced on episomal DNA, for example on T-DNA during transient transformation by *A. tumefaciens*. To analyze this possibility we generated three different sgRNA cleavage reporters based on a T-DNA containing a silent GUS reporter (Figure 6A). All three reporters contain a GUS reporter gene fused to minimal synthetic promoters that possess only low basal activity. Among them is the minimal 35S promoter fragment similar to the promoter that is fused to dTALE_BETA_ in the PLTB-locus. In addition, we chose a synthetic dTALE-Activated Promoter (STAP) and a minimal Bs4 promoter, because it was shown in previous studies that transient expression of these promoter scaffolds does not produce significant background GUS-activity (42,43). The target sequence of the tested sgRNA B1 was inserted upstream of these promoters at different distances to the ATG of the GUS reporter (Figure 6A). Remarkably, all reporters led to increased GUS activity compared to a negative control if Cas9 is co-expressed with the corresponding sgRNA B1 (Figure 6B). The STAP-based GUS reporter gave the strongest fold change (Figure 6C). These data show that a DSB on a T-DNA can also lead to activation of transcription and the production of translatable dilRNAs. Thus, DSBs, regardless of their location in the nucleus (genomic or episomal) lead to the production of transcripts, which can then be translated to a protein. This finding also shows that T-DNA-based reporters can be used to monitor sgRNA cleavage activity by detection of dilRNAs independently of the repair outcome, as shown with out-of-frame GUS reporters (44).

**Figure 6.**
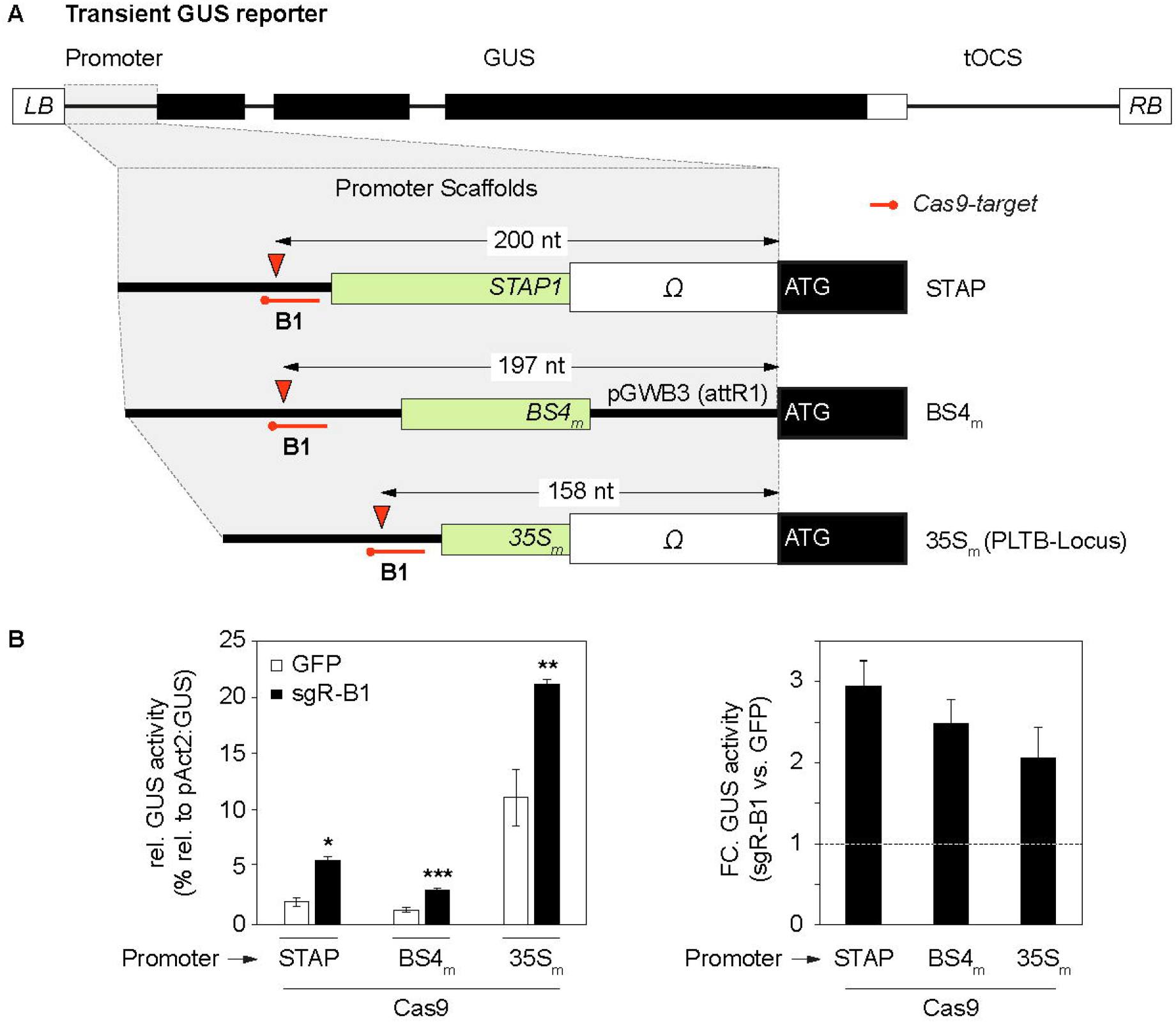
Production of dilRNAs can be monitored by transient reporters. **(A)** Schematic overview of tested synthetic GUS reporters. All GUS-fused promoters can be cleaved by sgRNA B1. Promoter elements and enhancer elements are indicated in green and white, respectively. The distance between the cleavage site and the GUS start codon is given. **(B)** Quantitative GUS measurement relative to Actin2 promoter-driven GUS. Samples have been harvested 3 dpi after transient expression of the indicated constructs. Combination of Cas9 with GFP serves as negative control. (Student’s t-test; *P-value ≤ 0.05; **P-value ≤ 0.01; ***P-value ≤ 0.001) **(C)** Fold change of GUS activity (Cut vs. uncut) with individual reporters.

## DISCUSSION

### Damage-induced RNAs can be translated

The presence of a silent transgene encoding a promoterless *dTALE* in the PTLB transgenic construct allows the detection of small increases in transcription thanks to the amplification of expression of reporter genes that are targets of the dTALE_BETA_. Using this setup we could show that DSBs lead to on target *de novo* transcription of dilRNAs, which could also be used for translation as *bona fide* mRNAs. The fact that all active sgRNAs tested in this study led to dilRNA formation and that dilRNA could be induced in transgenes, endogenes and extrachromosomal T-DNAs indicates that the production of dilRNAs upon DNA cleavage is a default and systematic process (Figure 2 and Figure 4). According to our data the level of dilRNAs represents 3-5% of the transcripts induced by dCas9-TV (Figure 4C, compare sgR-B1 and B2), indicating that the level of transcription induced by DSB is low, probably explaining why this phenomenon was not observed previously. One reason for the low levels of transcription could be that once repair of the DSB by error-prone NHEJ has occurred, Cas9 will not be able to cleave the DNA any longer. Thus, transcription could only take place either as long as the DSB is not repaired, or if there is repeated cleavage in the case of error-free repair. The fact that we were able to clearly detect dilRNA only on the side of the DSB that has a full transcription unit with a transcription terminator, supports the model that dilRNAs become *bona fide* mRNA transcripts when the genomic context allows it (Figure 2, 4C and 5A, Figure 7). Our findings demonstrate that a DSB in the neighborhood of a silent gene can lead to its activation. DSBs can occur more or less randomly upon genotoxic stress, for instance upon exposure to DNA damaging radiation. Therefore, since a relatively large fraction of plant genomes is made up of transposable elements (TEs; or TE leftovers; Arabidopsis 20%, Maize 80%; (45)), it is likely that some DSBs will occur upstream of TEs, thereby induce their transcription and possibly trigger transposition. DSB-induced TE transposition will generate genome rearrangements and could thus contribute to adaptation to specific environments if said rearrangements confer a selective advantage. DSBs could also occur in a targeted fashion and induce transcription as part of a developmental or physiological program. The only example we could find in the literature is in mouse, where neuronal activity triggers the targeted formation of DSBs in the vicinity of early response genes, whose transcription is then induced (46). In this case, the DSBs appear to be generated by the activity of topoisomerase II, which is preferentially bound to the upstream region of the targeted genes. Whether such targeted DSBs inducing transcription also occur in plants is currently unknown.

**Figure 7.**
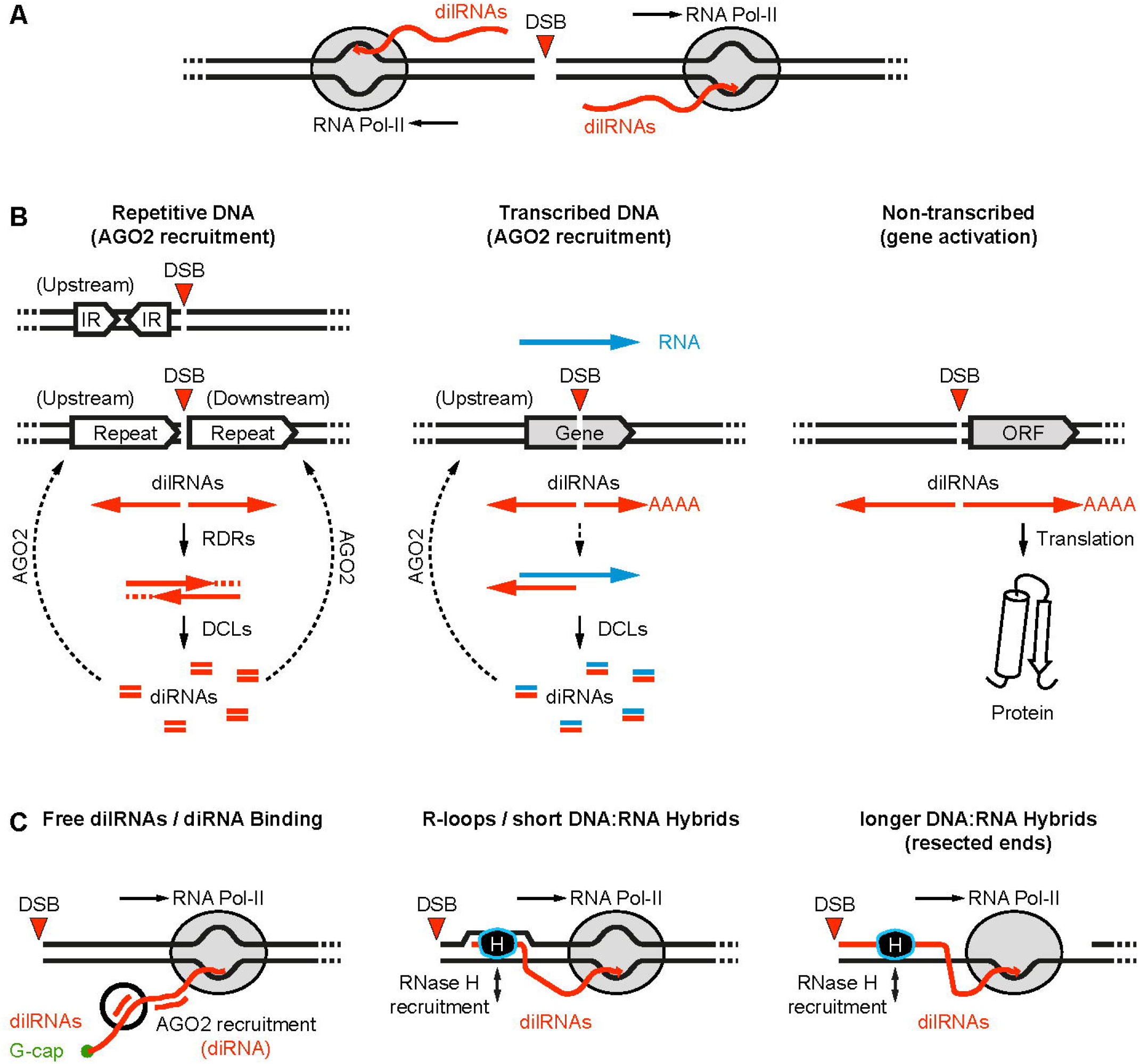
Hypothetical model of context dependent signaling activities by dilRNAs. **(A)** DSBs in plants led to de novo transcription of divergent dilRNAs. **(B)** Hypothetical context dependent signaling activities of dilRNAs. DSBs in repetitive sequences led to the production of complementary dilRNAs. Complementary dilRNAs can anneal to each other forming partial dsRNAs which might be extended by RDRs and further processed by DCL Ribonucleases forming small diRNAs. Small diRNAs can be incorporated into AGO2, priming it for sequence specific binding to dilRNAs or DNA and consequent repair factor recruitment in the vicinity of the DSB. DSBs in the vicinity of inverted repeats would lead to dilRNAs that could form dsRNA with itself and being extended by RDRs, followed by diRNA signaling. Similar signaling can take place if DSBs occur in highly expressed loci. Here, dilRNAs can anneal to pre-existing complementary transcripts. Corresponding dsRNA-derived small diRNAs can guide AGO2 (and repair factors) only to one site of the lesion. If DSBs occur close by an open reading frame (ORF) the corresponding dilRNAs can be translated to proteins. **(C)** DilRNAs can be targeted by small diRNAs (with or without AGO2). DilRNAs could form short DNA:RNA hybrids (R-loops) with the template strand, displacing the non-template strand. On 5’ resected ends longer stretches of DNA:RNA hybrids can form with the single stranded 3’ overhang. DNA:RNA hybrids could protect the single stranded 3’ overhang from degradation or could serve as DSB-specific binding platform for the repair machinery (for instance via RNase H enzymes).

### Possible functions of dilRNAs and diRNAs in plants

In plants, relatively less is known about dilRNAs as well as small diRNAs. To the best of our knowledge, dilRNAs have not been described in plants before. However, according to our data, dilRNAs are likely to be always produced if a DSB occurs. Besides their destination as *bona fide* mRNAs leading to translation of neighboring genes, dilRNAs could also serve as signal to recruit factors for DSB repair (Figure 7). This could be accomplished via mRNA decapping enzymes or diRNA-loaded AGO2 mediated sequence specific dilRNA binding (6,47). dilRNAs could also anneal to their template DNA strand and form DNA:RNA hybrids. Recently, the role of DNA:RNA hybrids in DNA repair has received increased attention. Although no data is available from plants, the impact of DNA:RNA hybrids on DNA DSB repair was investigated in yeast and human cells. Notably, formation of DNA:RNA hybrids was detected in the vicinity of DNA DSBs and associated with DDR-signaling (5,6,8-10,19). However, the translation of dilRNAs that we observed implies that they are exported to the cytosol and thus would not be able to form DNA:RNA hybrids at the DSB site. To investigate potential roles of dilRNAs in DSB repair further studies need to be done, in particular to monitor the localization of dilRNAs as they are made.

In Arabidopsis, two studies investigated the role of small diRNAs in DSB repair using DGU.US or DU.GUS reporter systems, which allow monitoring of DSB repair respectively via the single strand annealing (SSA) or the synthesis dependent strand annealing (SDSA) pathways (14,16). Although both studies report the detection of small RNAs from the regions surrounding the DSB, they do not identify dilRNAs associated with these diRNAs. Our RNA sequencing analysis showed that if there is no complementary RNA in the vicinity of the DSB, hardly any small RNAs (diRNAs) can be detected. To the best of our knowledge, in all cases reported so far, small diRNAs are present only when the DSB occurs within an actively transcribed gene (14,16). Furthermore, our data show that long RNAs are produced upon DSB and therefore suggest that the small diRNAs that occur in actively transcribed genes derive from dsRNAs that form by annealing of complementary transcripts (dilRNA-dilRNAs; dilRNAs-mRNAs; Figure 7B). All other factors described to be involved in diRNA formation so far are most likely downstream of dilRNAs production and the annealing of complementary transcripts. For instance, by extension and amplification of complementary dilRNAs by RNA-dependent POLYMERASE 6 (RDR6), subsequent processing of the corresponding dsRNA into diRNAs by DICER-LIKE2-4 (DCL2-4) and their protection from degradation by incorporation of ARGONAUTE2 (AGO2) could occur (14). The observed positive effect of diRNAs on single strand annealing (SSA; (14)), a form of HDR, could not be confirmed by Miki and colleagues (16). Therefore, in plants, the role of diRNAs in the repair of DSBs is still an open question. However, the fact that diRNAs are stabilized by AGO2, and that AGO2 is responsible for RAD51 recruitment to DSBs, suggests a participation of diRNAs in DDR-signaling in plants (19). An open question is whether the presence of diRNAs can serve as a signal to modulate DDR locally, for example by promoting precise repair of highly expressed, essential housekeeping genes (Figure 7B).

### Which RNA polymerase generates DSB-induced long RNAs?

According to Wei and colleagues (14) the formation of diRNAs depends on RNA Pol-IV (NPRD1), whereas diRNA-formation is promoted by the deletion of RNA Pol-V (NRPE1). The question arises whether dilRNAs (precursor of diRNAs) are synthesized by RNA Pol-IV. RNA polymerases IV and V are plant-specific and participate in RNA-directed DNA methylation (RdDM) and possess changes in invariant or conserved residues compared to RNA Pol I-III, which underpins their specialized function (48,49). From a mechanistic point of view, RNA Pol-IV, compared to RNA Pol II, was described as a weak RNA polymerase requiring a RNA primer for elongation that is furthermore terminated if the polymerase encounters 12-18 nt of dsDNA (50). In addition, even with ssDNA as substrate, RNA Pol-IV was shown to generate only short RNA fragments (26-45 nt) and terminates transcription at methylated cytosines (51). Notably, the dilRNAs we identified are approx. 4 kb long (dTALE_BETA_), poly-adenylated and translated. Therefore, our data rather support that dilRNAs are produced by RNA Pol-II, as shown previously in mammalian cells (10,11).

### Model of DSB-induced transcription

In mammalian cells and yeast, it was shown that dilRNAs are produced from each side of DSBs, a phenomenon which, according to our data, is likely to occur in plants (Figure 5 and Figure 7A). We propose a general model in which transcription by RNA Pol II on either side of the DSB is one of the earliest events following DNA cleavage. This transcription generates long RNAs (dilRNAs) whose fate depends on the location of the DSB in the genome. If the DSB occurs upstream of a gene, for example in the promoter region, a stable dilRNA (mRNA) will be produced leading to the production of a protein. When the gene is silent, this can lead to de novo protein production and initiation of a signaling cascade if the gene codes for a transcription factor, or genome rearrangements if there is a transposable element. If the DSB occurs within an actively transcribed gene, then the antisense transcript induced by the DSB (dilRNA) can anneal to the sense transcript of the gene, triggering processing of the RNAs by DCLs and prime AGO proteins for sequence specific nucleic acid (diRNA/DNA, diRNA/dilRNA or diRNA/mRNA) binding. The dilRNA in the sense orientation can either be translated leading to the production of a truncated protein if there is an in frame ATG, or degraded if the there is no in frame ATG or a premature stop codon. In some cases, the presence of repeats, either inverted if they are on the same side of the DSB or direct if they are on either side, can lead to self-complementary or complementary RNAs which again can be processed by the DCL machinery. All these processes are short-lived, since once the DSB is repaired, DSB induced transcription will stop. The question therefore is whether DSB-induced transcription plays a role in the repair process as discussed above or if it is just a product of the affinity of RNA polymerase for broken DNA ends.

Our findings show that transcription of long RNAs is a default and early event in the processing of DSBs raising multiple questions with connections to the impact on DNA repair processes as well as on the role of DSBs in regulating gene expression.

## DATA AVAILIBILITY

All annotated sequences and used primers are available in the supporting information. The RNAseq data has been depositied in NCBI as Bioproject SUB11197124.

## ACKNOWLEDGEMENTS

We thank David Chiasson for the SlU6 promoter construct. We also thank the IPB gardeners and staff for excellent assistance.

## SUPPLEMENTARY FIGURES

**Supplementary Figure 1. (A)** Schematic presentation of the pAGM26035 transgene in *Nicotiana benthamiana* (Nb) primary transformants (pt). The presence of the functional transgene can be confirmed by transient expression of dTALE_BETA_ and resulting upregulation of Betalain biosynthesis genes (5GT, DODA1, Cyp76AD1). **(B)** Photos of inoculated plants 4 dpi. Red coloration confirms presence of the transgene. **(C)** Identification of T-DNA insertion position in the Nb genome. T-DNA flanking gDNA regions could be identified for pt12 and pt57. PCR with corresponding primers using genomic DNA (gDNA) as template confirmed the point of T-DNA insertion. pAGM26035 pt57 and pt 81 might have been derived from the same callus. Nb pAGM26035 pt12 and pt57 have been used for further characterization.

**Supplementary Figure 2. Confirmation of the genomic T-DNA-insertion points and functional validation of the TALE-Betalain circuit in transgenic *Nicotiana benthamiana* (Nb) line pAGM25036 primary transformant pt12 (A) and pt57 (B)**. Schematic presentation of dTALE-and Cas9-target sites in the T-DNA-flanking genomic DNA for each transformant is given in the upper panel. Pictures were taken four and seven days post inoculation (dpi) of Agrobacterium strains, leading to expression of the indicated constructs. Red coloration is a result of Betalain biosynthesis initiated by the activated dTALE_BETA_ transgene. Transient expression of dTALE_BETA_ (35S:dTALE_BETA_) and GFP (35S:GFP) served as positive and negative control, respectively.

**Supplementary Figure 3. Analysis of sgRNA B4 cleavage-activity by amplified fragment length polymorphism (ALFP). (A)** Overview of used sgRNAs and primers for AFLP. Corresponding constructs have been transiently expressed in Nb PLTB pt57-line via Agrobacterium. **(B)** sgRNA T1 mediates low levels of target DNA cleavage. Four dpi gDNA was isolated and used for PCR using primers F2 and R. If both sgRNAs are active, NHEJ-mediated repair of the Cas9 induced DNA double strand breaks led to a detectable deletion. No deletion could be observed if sgRNA B3 is combined with sgRNA T1.

**Supplementary Figure 4. Cas9 nickases do not induce dilRNAs (A)** Schematic overview of tested Cas9 target sites in the PLTB-locus. **(B)** Quantitative GUS measurement after transient expression of Cas9-variants together with corresponding sgRNAs (one sgRNA) and STAP1_BETA_:GUS reporter. DNA strands which could be used by the RNA polymerase as substrate for transcription of dTALE_BETA_ is indicated in green. Cas9 WT cleaves both strands, Cas9(D10A) and Cas9(H840A) cleave the targeted and non-targeted strand only, respectively. Single guide RNA D1 (non-targeting control) and dCas9 serve as negative controls. **(C)** Quantitative GUS measurement after transient expression of Cas9-variants together with corresponding sgRNAs (dual sgRNAs) and STAP1_BETA_:GUS reporter. Cleavage sites of sgR-B1 and sgR-B3 are 68 nt apart. Combination of dual sgRNAs with Cas9(D10A) and Cas9(H840A) lead to 68 nt 5’ and 68 nt 3’ overhangs, respectively. (Student’s t-test; *P-value ≤ 0.05; **P-value ≤ 0.01; ***P-value ≤ 0.001)

**Supplementary Figure 5. Cas12a mediated DSBs induce dilRNAs (A)** Schematic overview of tested Cas9- and Cas12a target sites in the PLTB-locus. **(B)** Quantitative GUS measurement after transient expression of Cas9- or Cas12a-variants together with corresponding sgRNAs (Cas9) or crRNAs (Cas12a) and STAP1_BETA_:GUS reporter. Cleavage patterns of Cas9 and Cas12a are given on the right panel. Cas9 mainly produces blunt ends (3 nt upstream of PAM) and Cas12a mainly short 5’ sticky ends (7 nt overhang). Deactivated Cas12a (dCas12a) and engineered dCas12a-based transcriptional activator (dCas12a-TV) served as negative and positive control, respectively. (Student’s t-test; *P-value ≤ 0.05; **P-value ≤ 0.01; ***P-value ≤ 0.001)

**Supplementary Figure 6. GUS assay parallel to RNA isolation for RT-PCR (figure 4). (A)** Schematic overview of the PLTB-locus. Used Cas9 sgRNA target sites are indicated **(B)** Quantitative GUS measurement after transient expression of indicated constructs together with the STAP1BETA:GUS reporter. Values are relative to GUS activity of Cas9 without sgRNA. Note that samples for the GUS assay and RNA isolation were harvested from the same inoculation spot but GUS assay was only performed with two individual plants. (Student’s t-test; *P-value ≤ 0.05; **P-value ≤ 0.01; ***P-value ≤ 0.001)

**Supplementary Figure 7. (A)** Agarose gel showing nested PCR products from G-tailed cDNA (oligo dT) from indicated samples. Left planel shows the result of the first experiment comparing activator-induced transcripts (dCas9-TV, dTALE(pt57-2)) against DSB-induced transcripts (Cas9 with sgRNA G3, B1 and B3). Right panel showed second experiment with activators only. **(B)** Sequencing results of nested PCR products from G-tailed cDNA (oligo dT). Length correspond to the position within the PLTB-transgene. Number indicates number of reads.

**Supplementary Figure 8. (A)** A primer set for the Nb PLTB pt57 transgene locus was tested by PCR using genomic DNA as template. Note that the PCR master mix contains the gDNA template and an access of specific primers was added to each reaction. Fragment 2 and fragment 4 could only be amplified with lower amounts. Primers used for RT-PCR are indicated in red. **(B)** Oligo-dT synthesized cDNAs, which have been used for RT-PCR (Figure 4B and C) were analysed for gDNA contamination by PCR. Control primers that bind to endogenous intronic regions could only led to an amplicon in the presense of gDNA. No gDNA contamination could be observed for used cDNAs. Genomic DNA from Nb PLTB pt57 and Nb WT served as positive controls.

**Supplementary Figure 9**. GUS assay parallel to RNA isolation for RNA seq (figure 5). (A) Schematic overview of the PLTB-locus. The target site of sgRNA B1 is indicated (B) Quantitative GUS measurement after transient expression of sgRNA B1 with or without Cas9 together with the STAP1_BETA_:GUS reporter. Values are relative to pAct2:GUS activity. Note that samples for the GUS assay and RNA isolation were harvested from the same inoculation spot. (Student’s t-test; *P-value ≤ 0.05; **P-value ≤ 0.01; ***P-value ≤ 0.001)**Supplementary Figure 10. Mapped small RNA reads originated from transiently expressed T-DNAs. (A)** Sequence alignments of transiently expressed T-DNA and the PLTB transgene. Sequences and homologous parts are indicated. **(B)** Small RNA reads mapped to the PLTB transgene derived from “Cas9” sample. Origin of small reads gets visible at the borders of homology. Read coverage summary (from Figure 5) is given in the lower left panel. **(C)** Small RNA reads mapped to the PLTB transgene derived from “no Cas9” sample. Origin of small reads gets visible at the borders of homology. Read coverage summary (from Figure 5) is given in the lower left panel.

**Supplementary Figure 10**. Mapped small RNA reads originated from transiently expressed T-DNAs. (A) Sequence alignments of transiently expressed T-DNA and the PLTB transgene. Sequences and homologous parts are indicated. (B) Small RNA reads mapped to the PLTB transgene derived from “Cas9” sample. Origin of small reads gets visible at the borders of homology. Read coverage summary (from figure 5) is given in the lower left panel. (C) Small RNA reads mapped to the PLTB transgene derived from “no Cas9” sample. Origin of small reads gets visible at the borders of homology. Read coverage summary (from figure 5) is given in the lower left panel.

