## Supplemental Figures for "DNA double strand breaks lead to *de novo* transcription and translation of damage-induced long RNAs *in planta*"

#### Supplemental Figure 1

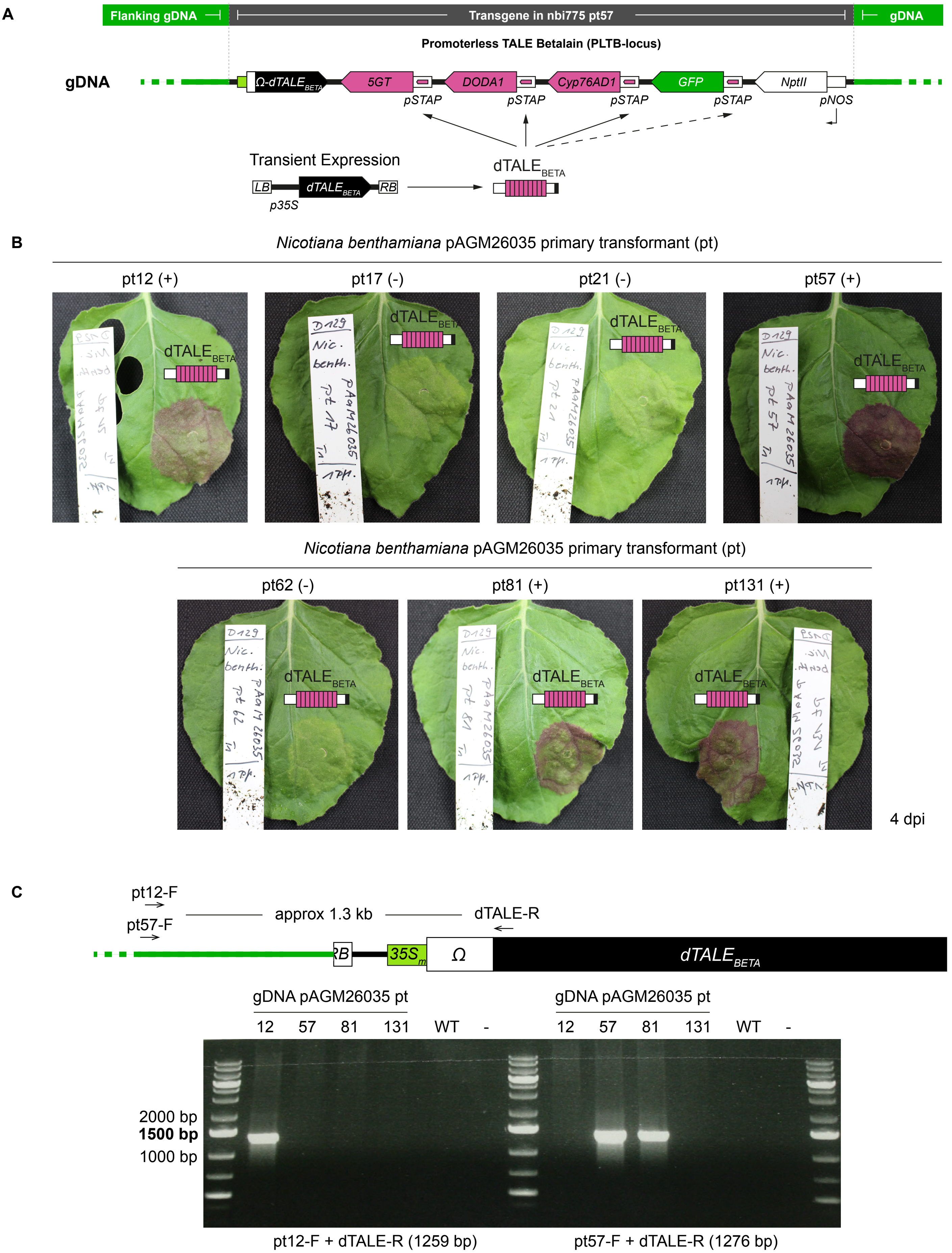


**Supplementary Figure 1. (A)** Schematic presentation of the pAGM26035 transgene in Nicotiana benthamiana (Nb) primary transformants (pt). The presence of the functional transgene can be confirmed by transient expression of dTALE$_{BETA}$ and resulting upregulation of Betalain biosynthesis genes (5GT, DODA1, Cyp76AD1). **(B)** Photos of inoculated plants 4 dpi. Red coloration confirms presense of the transgene. **(C)** Identification of T-DNA insertion position in the Nb genome. T-DNA flanking gDNA regions could be identified for pt12 and pt57. PCR with corresponding primers using genomic DNA (gDNA) as template confirmed point of T-DNA insertion. pAGM26035 pt57 and pt 81 might have been derifed from the same callus. Nb pAGM26035 pt12 and pt57 have been used for further characterization.

Supplemental figure 2

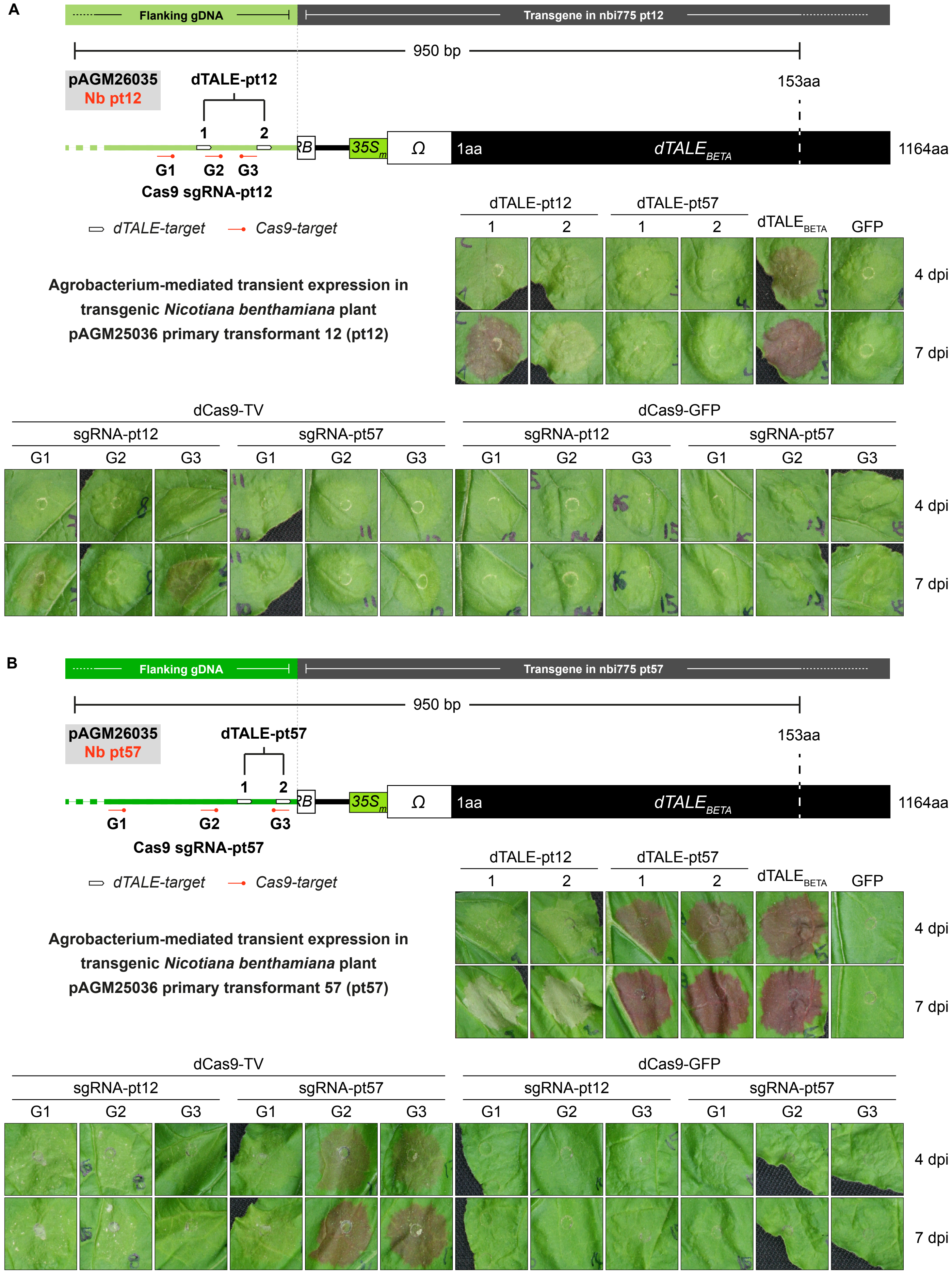


**Supplementary Figure 2. Confirmation of the genomic T-DNA-insertion points and functional validation of the TALE-Betalain circuit in transgenic *Nicotiana benthamiana* (Nb) line pAGM25036 primary transformant pt12 (A) and pt57 (B).** Schematic presentation of dTALE- and Cas9-target sites in the T-DNA-flanking genomic DNA for each transformant is given in the upper panel. Pictures were taken four and seven days post inoculation (dpi) of Agrobacterium strains, leading to expression of the indicated constructs. Red coloration is a result of Betalain biosynthesis initiated by the activated $dTALE_{BETA}$ transgene. Expression of $dTALE_{BETA}$ and GFP served as positive and negative control, respectively.

#### Supplementary Figure 3

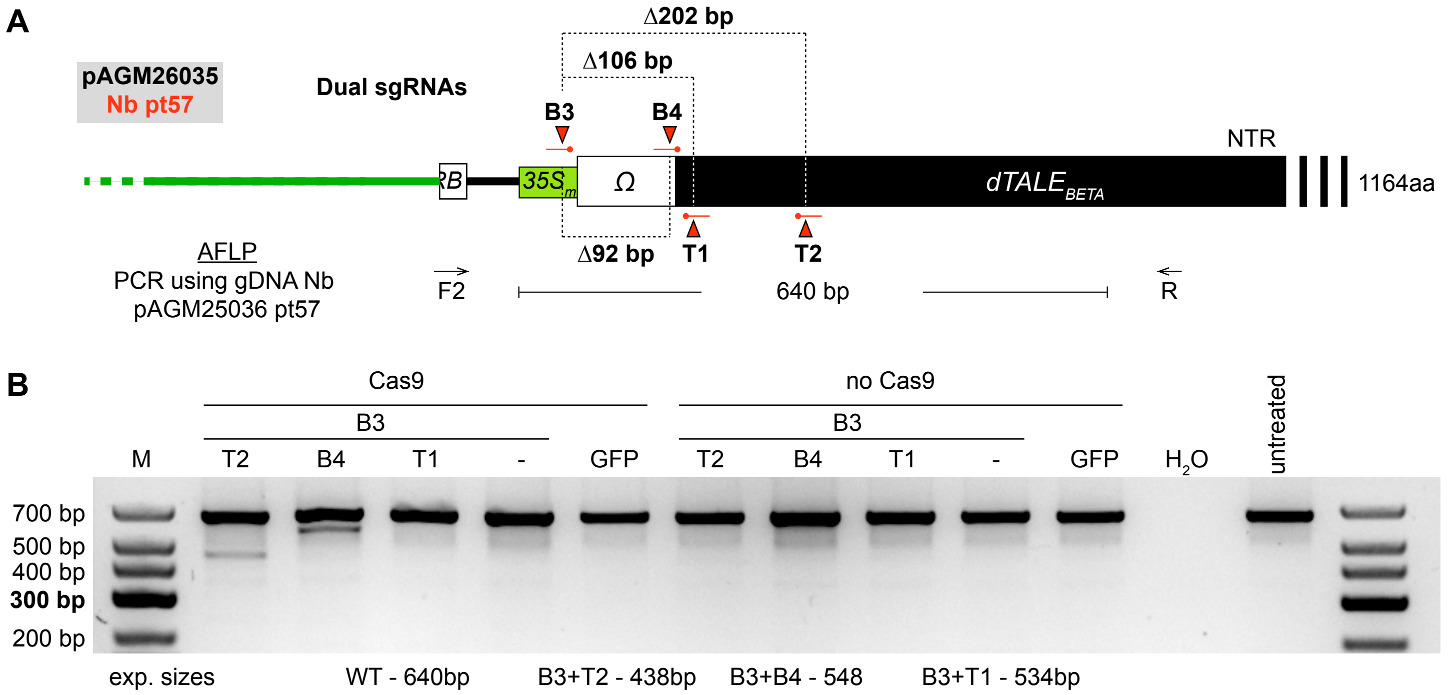


**Supplementary Figure 3. Analysis of sgRNA B4 cleavage-activity by amplified fragment length polymorphism (ALFP). (A)** Overview of used sgRNAs and primers for AFLP. Corresponding contructs have been transiently expressed in Nb PLTB pt57-line via Agrobacterium. **(B)** sgRNA T1 mediates low levels of target DNA cleavage. Four dpi gDNA was isolated and used for PCR using primers F2 and R. If both sgRNAs are active, NHEJ-mediated repair of the Cas9 induced DNA double strand breaks led to a detectable deletion. No deletion could be observed if sgRNA B3 is combined with sgRNA T1.

#### Supplementary Figure 4

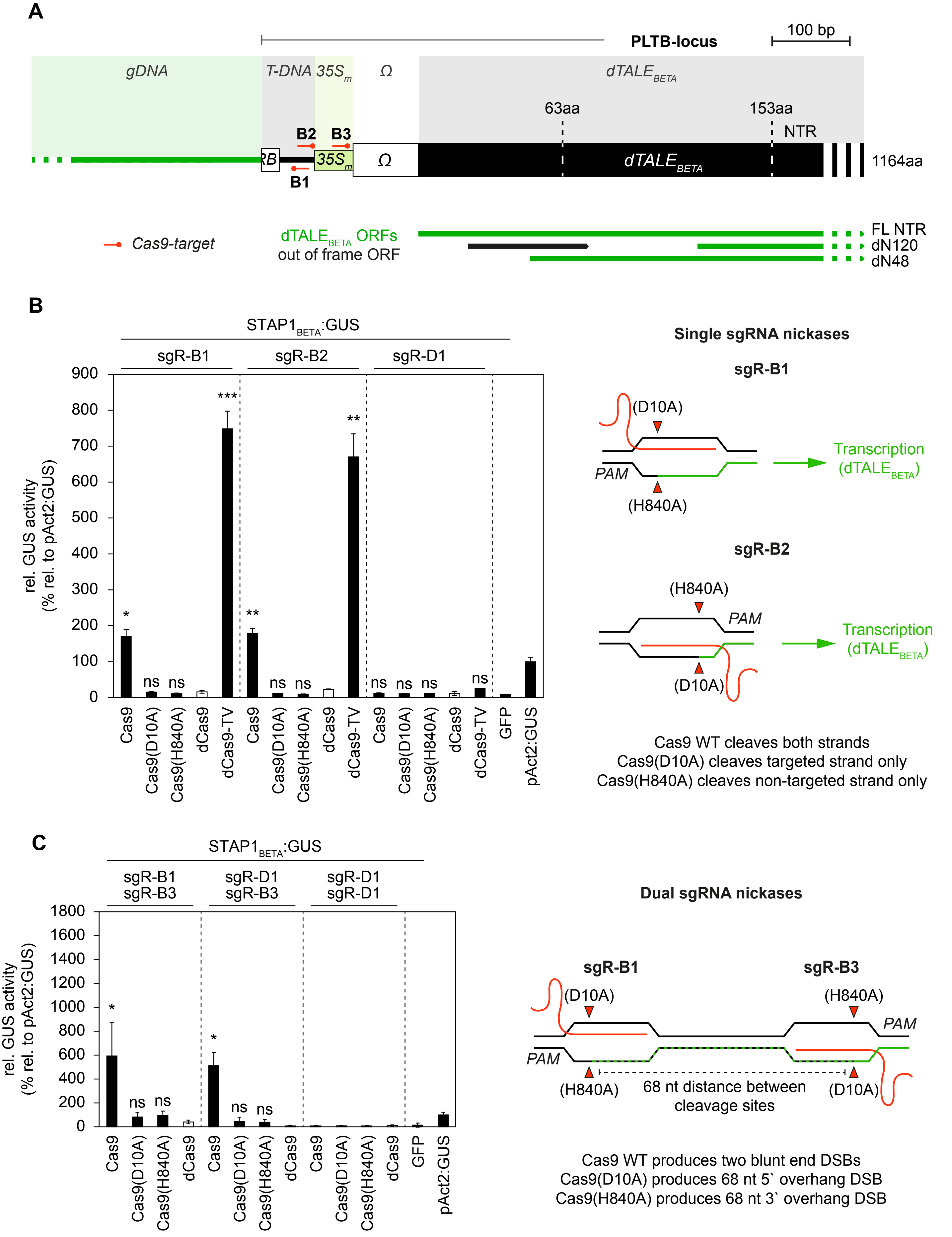


**Supplementary figure 4. Cas9 nickases do not induce diIRNAs (A)** Schematic overview of tested Cas9 target sites in the PLTB-locus. **(B)** Quantitative GUS measurement after transient expression of Cas9-variants together with corresponding sgRNAs (one sgRNA) and $STAP1_{BETA}$:GUS reporter. DNA strands which could be used by the RNA polymerase as substrate for transcription of $dTALE_{BETA}$ is indicated in green. Cas9 WT cleaves both strands, Cas9(D10A) and Cas9(H840A) cleave the targeted and non-targeted strand only, respectively. Single guide RNA D1 (non-targeting control) and dCas9 serve as negative controls. **(C)** Quantitative GUS measurement after transient expression of Cas9-variants together with corresponding sgRNAs (dual sgRNAs) and $STAP1_{BETA}$:GUS reporter. Cleavage sites of sgR-B1 and sgR-B3 are 68 nt apart. Combination of dual sgRNAs with Cas9(D10A) and Cas9(H840A) lead to 68 nt 5´ and 68 nt 3´ overhangs, respectively. (Student's t-test; *P-value ≤ 0.05; **P-value ≤ 0.01; ***P-value ≤ 0.001).

#### Supplementary Figure 5

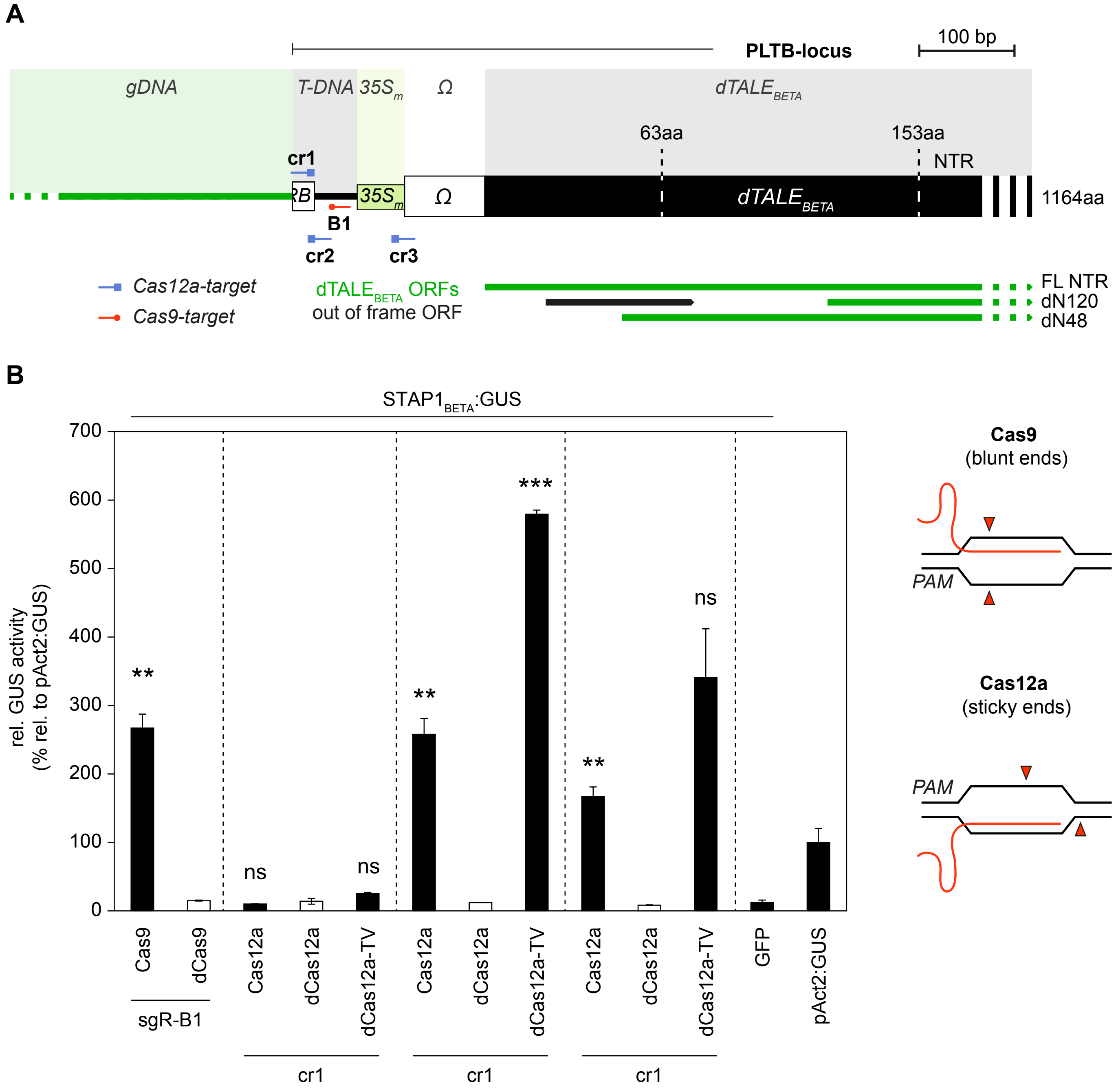


**Supplementary Figure 5. Cas12a mediated DSBs induce diIRNAs (A)** Schematic overview of tested Cas9- and Cas12a target sites in the PLTB-locus. **(B)** Quantitative GUS measurement after transient expression of Cas9- or Cas12a- variants together with corresponding sgRNAs (Cas9) or crRNAs (Cas12a) and $STAP1_{BETA}$:GUS reporter. Cleavage patterns of Cas9 and Cas12a are given on the right panel. Cas9 mainly produces blunt ends (3 nt upstream of PAM) and Cas12a mainly short 5´ sticky ends (7 nt overhang). Deactivated Cas12a (dCas12a) and engineered dCas12a-based transcriptional activator served as negative and positive control, respectively. (Student's t-test; *P-value ≤ 0.05; **P-value ≤ 0.01; ***P-value ≤ 0.001)

#### Supplemental Figure 6

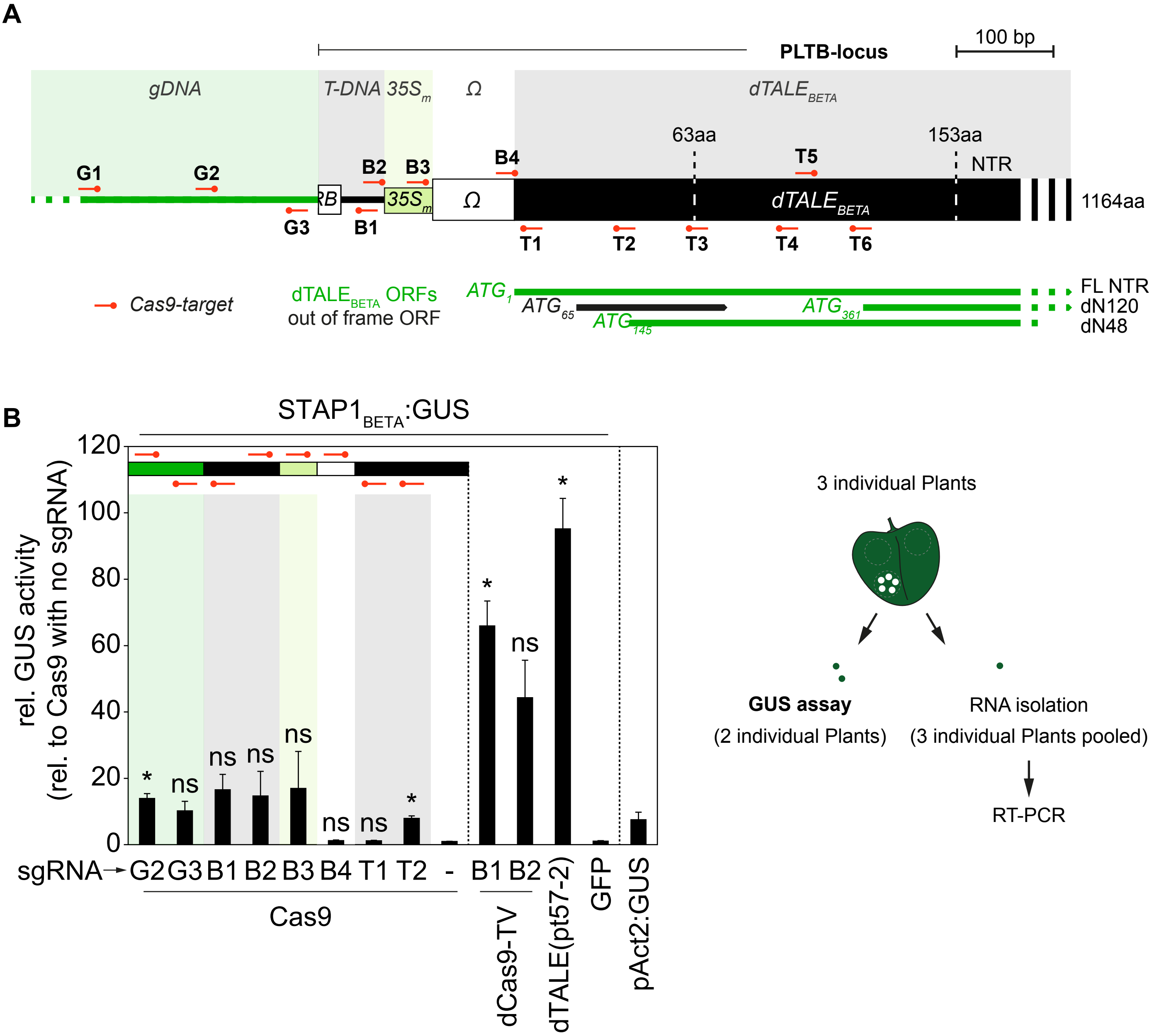


**Supplementary Figure 6. GUS assay parallel to RNA isolation for RT-PCR (figure 4).** **(A)** Schematic overview of the PLTB-locus. Used Cas9 sgRNA target sites are indicated **(B)** Quantitative GUS measurement after transient expression of indicated constructs together with the $STAP1_{BETA}$:GUS reporter. Values are relative to GUS activity of Cas9 without sgRNA. Note that samples for the GUS assay and RNA isolation were harvested from the same inoculation spot but GUS assay was only performed with two individual plants. (Student's t-test; *P-value ≤ 0.05; **P-value ≤ 0.01; ***P-value ≤ 0.001)

#### Supplementary Figure 7

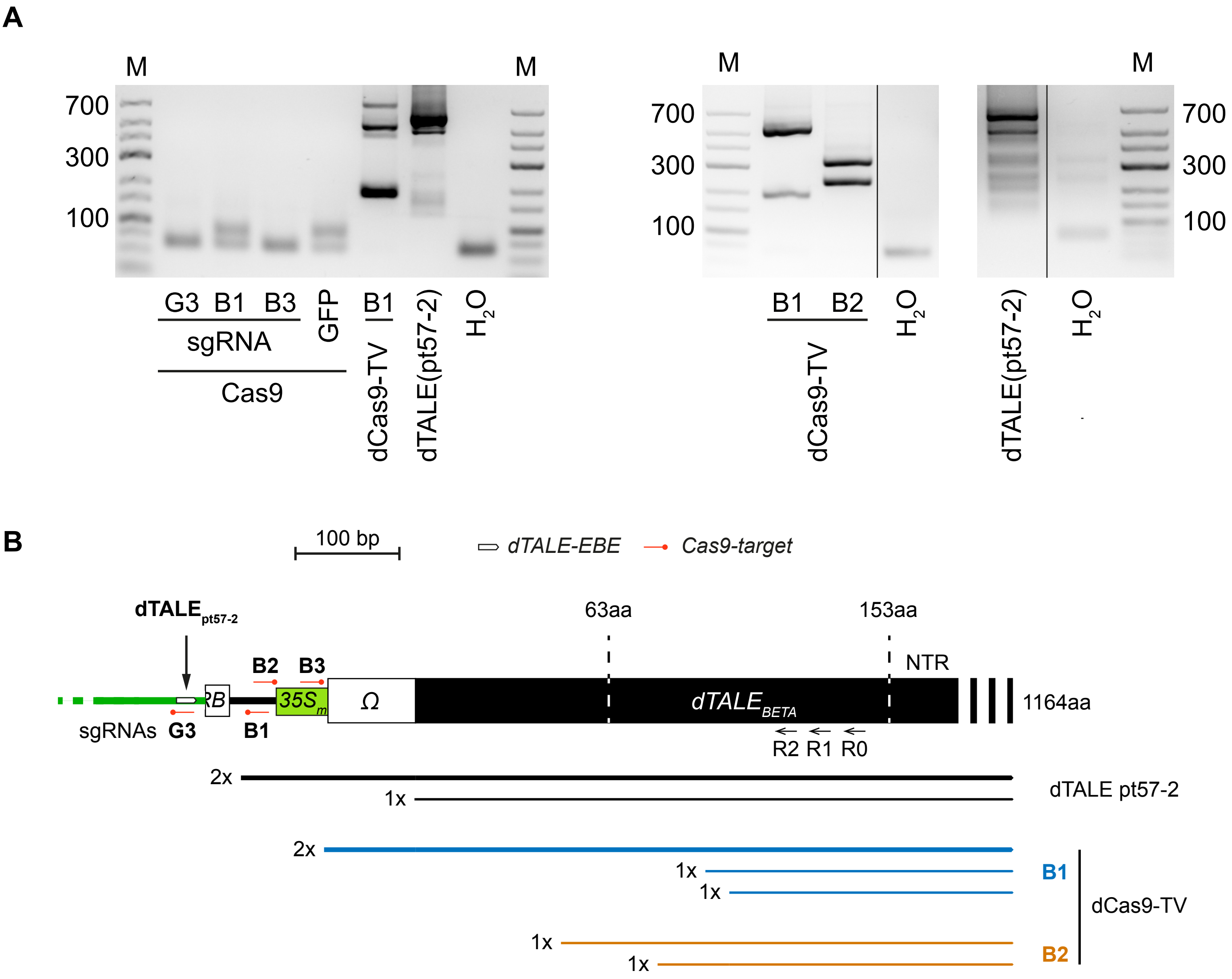


**Supplementary Figure 7. (A)** Agarose gel showing nested PCR products from G-tailed cDNA (oligo dT) from indicated samples. Left planel shows the result of the first experiment comparing activator-induced transcripts (dCas9-TV, dTALE(pt57-2)) against DSB-induced transcripts (Cas9 with sgRNA G3, B1 and B3). Right panel showed second experiment with activators only. **(B)** Sequencing results of nested PCR products from G-tailed cDNA (oligo dT). Length corrspond to the position within the PLTB-transgene. Number indicates number of reads.

#### Supplemental Figure S8

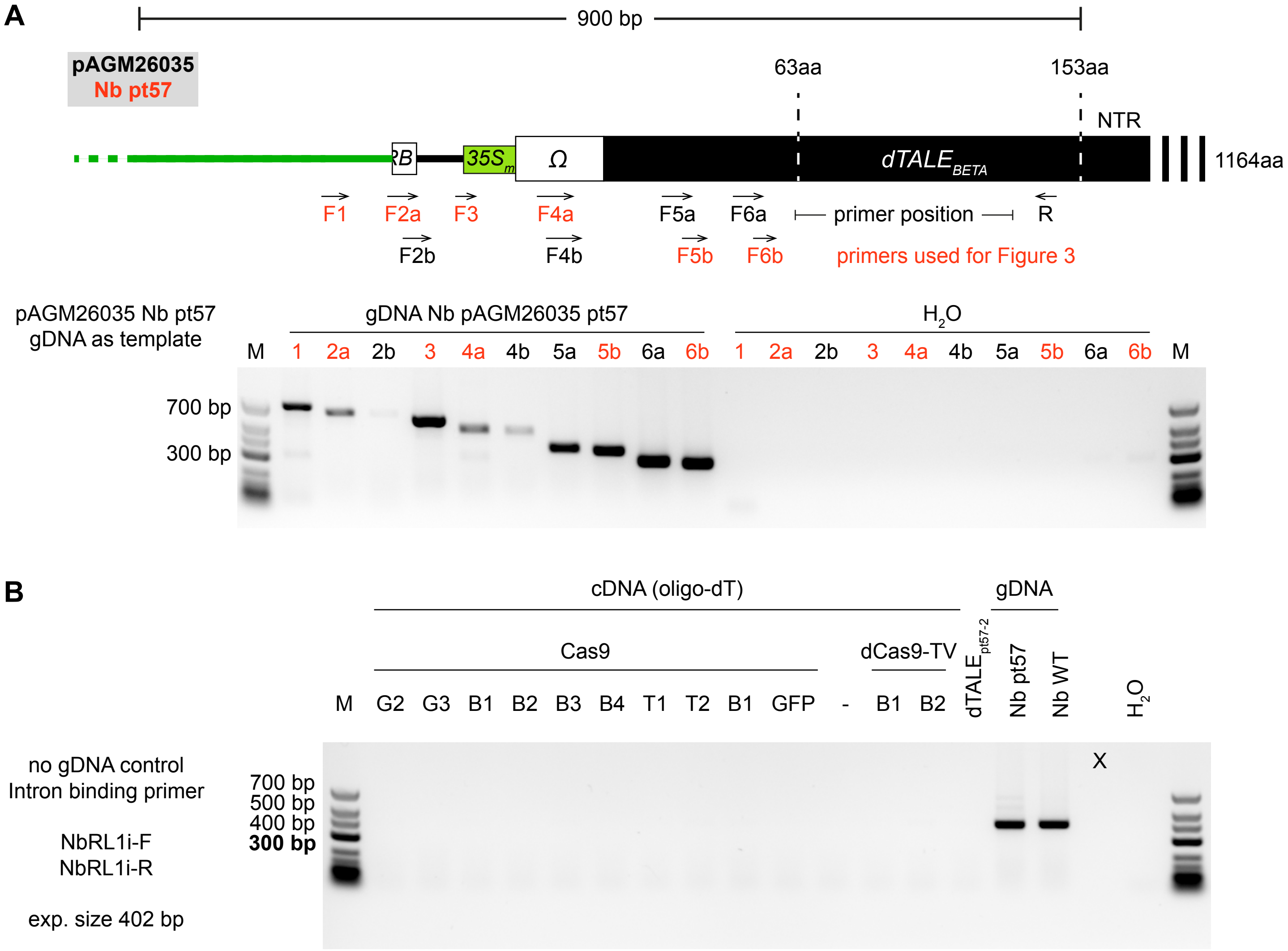


**Supplementary Figure 8. (A)** A primer set for the Nb pt57 transgene locus was tested by PCR using genomic DNA as template. Note that the PCR master mix contains the gDNA template and an access of specific primers was added to each reaction. Fragment 2 and fragment 4 could only be amplified with lower amounts. Primers used for RT-PCR are indicated in red. **(B)** Oligo-dT synthesized cDNAs, which have been used for pRT-PCR (Figure 4B and 4C) were analysed for gDNA contamination by PCR. Control primers that bind to endogenous intronic regions could only led to an amplicon in the presense of gDNA. No gDNA contamination could be observed. Genomic DNA from Nb PLTB pt57 and Nb WT served as positive controls.

**Supplementary Figure 9**

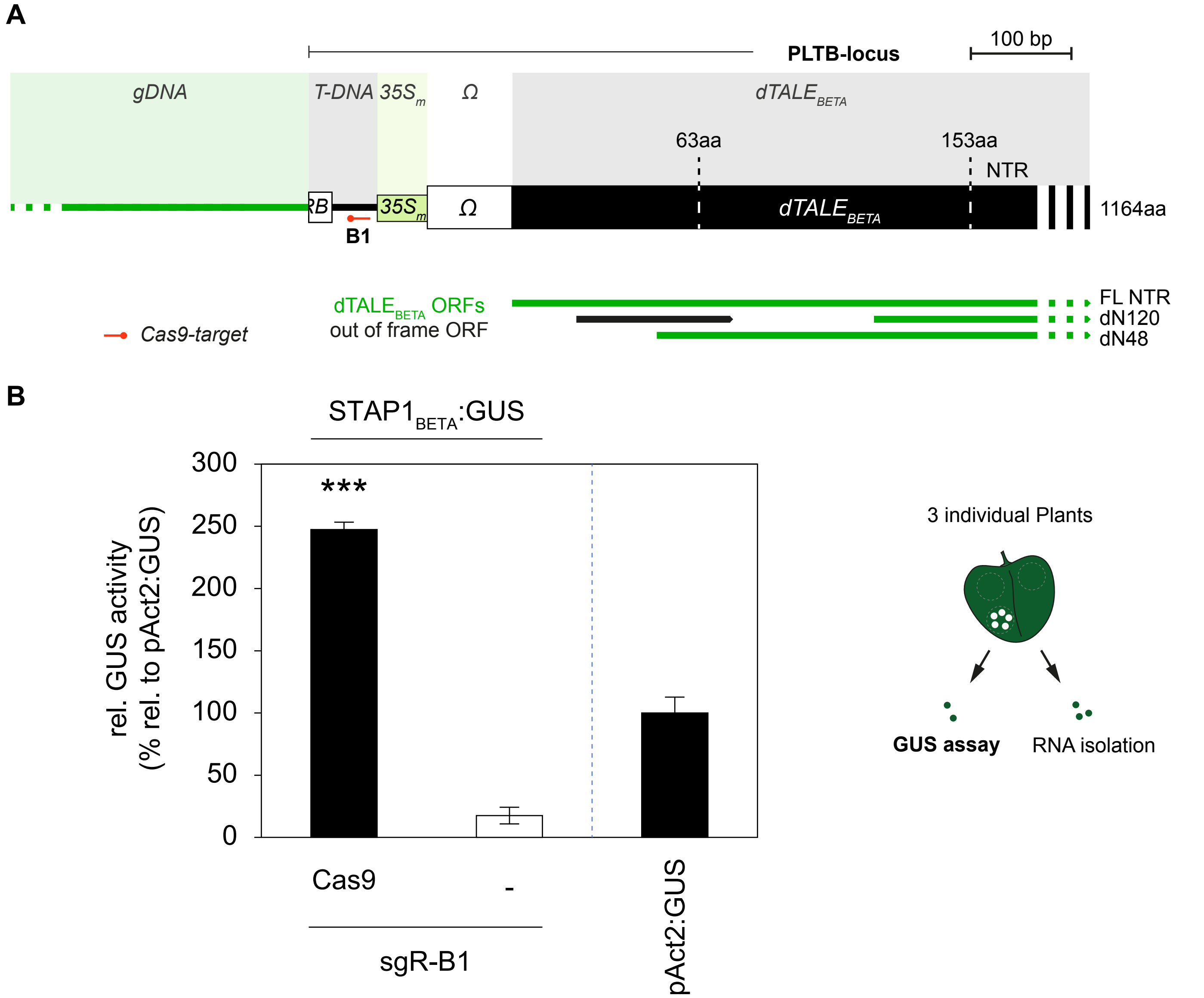


**Supplementary Figure 9. GUS assay parallel to RNA isolation for RNA seq (figure 5)**. (A) Schematic overview of the PLTB-locus. The target site of sgRNA B1 is indicated **(B)** Quantitative GUS measurement after transient expression of sgRNA B1 with or without Cas9 together with the $STAP1_{BETA}$:GUS reporter. Values are relative to pAct2:GUS activity. Note that samples for the GUS assay and RNA isolation were harvested from the same inoculation spot. (Student's t-test; *P-value ≤ 0.05; **P-value ≤ 0.01; ***P-value ≤ 0.001)

### Figure S10A – Sequence alignments of the transiently expressed T-DNAs (sgR-B1, Cas9, GUS-BAR) to the PLTB pt57 transgene

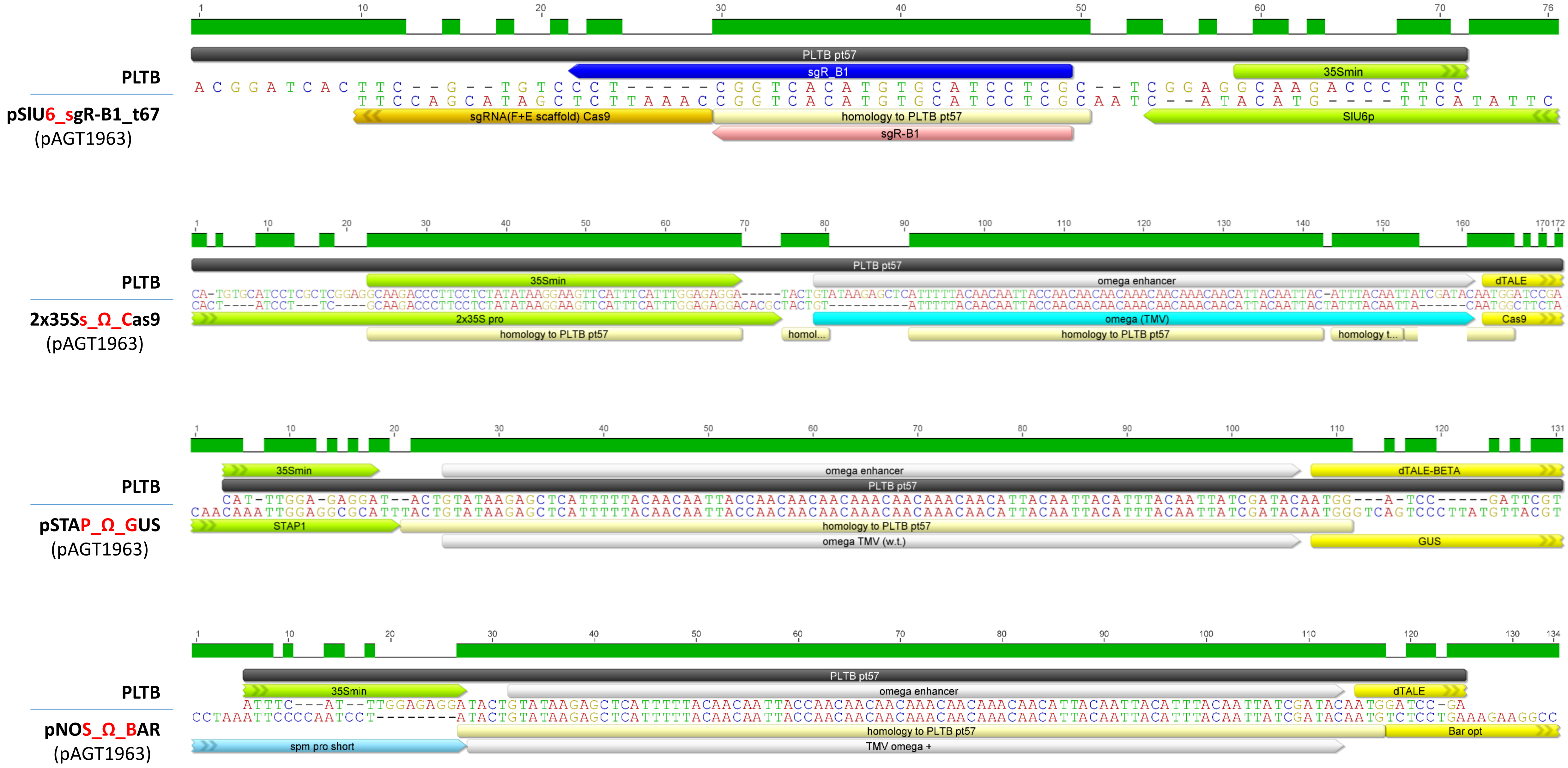

**Figure S10B - Small RNA reads from „Cas9“ sample mapped to PLTB pt57 transgene**

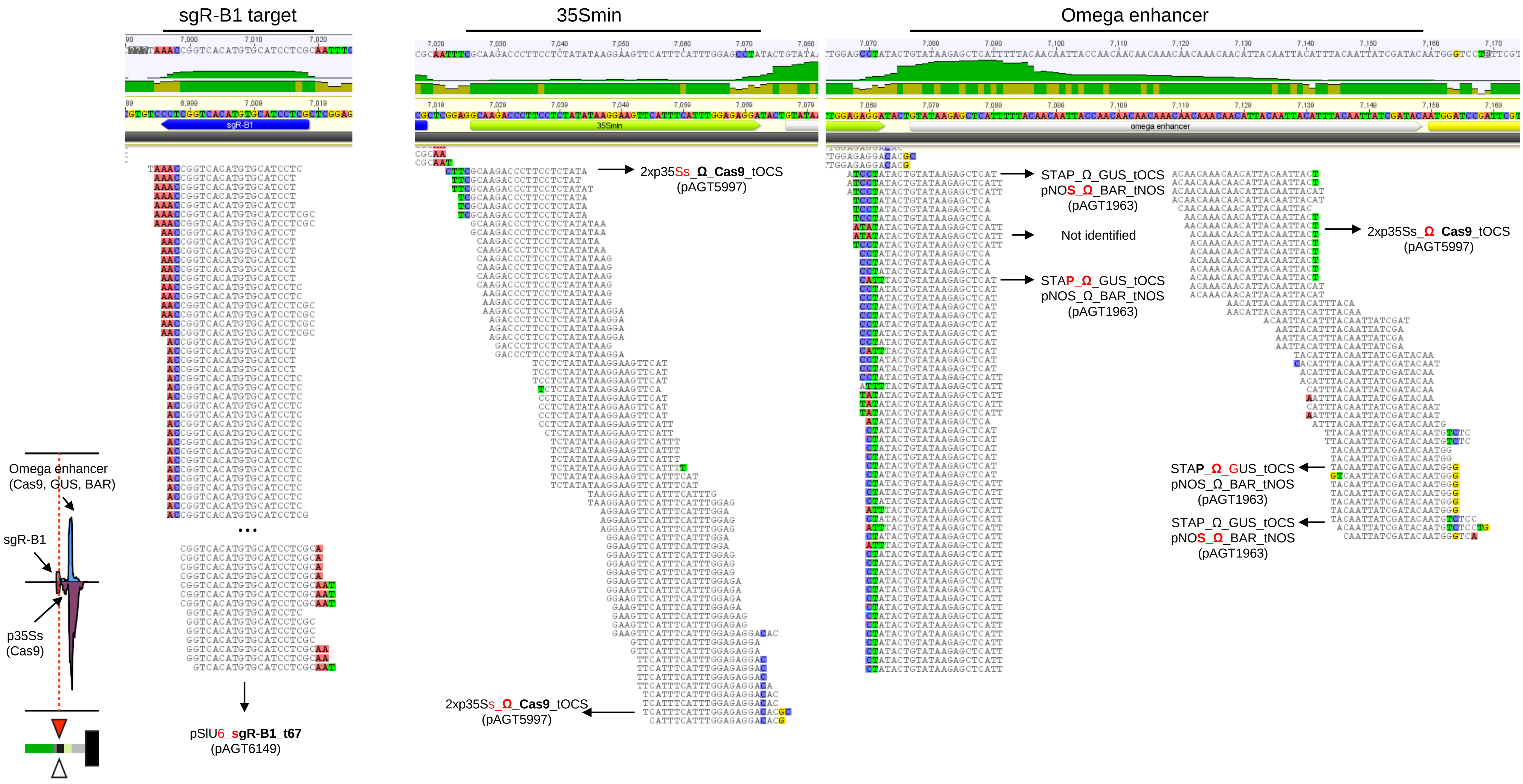

#### Figure S10C - Small RNA reads from „no Cas9“ sample mapped to PLTB pt57 transgene

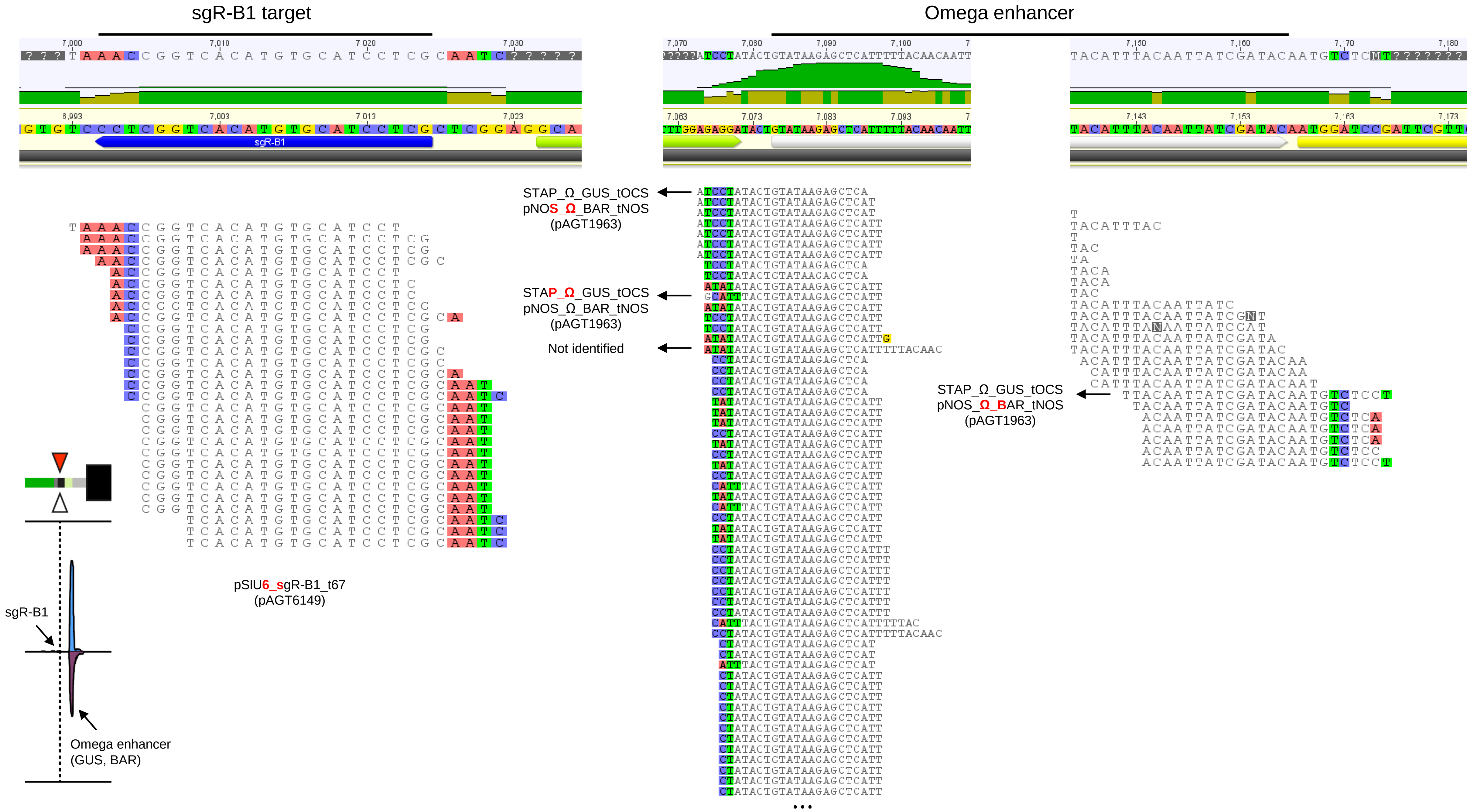

**Supplementary Figure 10. Mapped small RNA reads originated from transiently expressed T-DNAs. (A)** Sequence alignments of transiently expressed T-DNA and the PLTB transgene. Sequences and homologous parts are indicated. **(B)** Small RNA reads mapped to the PLTB transgene derived from “Cas9” sample. Origin of small reads gets visible at the borders of homology. Read coverage summary (from figure 5) is given in the lower left panel. **(C)** Small RNA reads mapped to the PLTB transgene derived from “no Cas9” sample. Origin of small reads gets visible at the borders of homology. Read coverage summary (from figure 5) is given in the lower left panel.
